# Prokaryotic Community Structures in the Bering and Chukchi Seas

**DOI:** 10.64898/2026.11.02.763580

**Authors:** Rei Shibue, R. Eric Collins

## Abstract

Bacteria are essential to marine ecosystems as they remineralize organic matter, cycle carbon, nitrogen, and other key elements, and support higher trophic levels through food webs. The Bering and Chukchi Seas off the coast of Alaska are among the most productive regions in the world, with intense seasonal algal blooms, yet their bacterial communities remain poorly characterized, and the region is warming rapidly as heat transport from the Pacific Ocean increases. Here, we used 16S rRNA gene amplicon sequencing of 181 samples collected on 0.2 µm filters at three depths (surface, middle, and bottom) across 62 stations during the Arctic Shelf Growth, Advection, Respiration and Deposition Rate Experiments (ASGARD) cruise in June 2017, when sea surface temperatures were anomalously warm. Hierarchical clustering of amplicon sequence variant (ASV) relative abundances resolved 11 sample clusters in three divisions, all dominated by Alphaproteobacteria, Gammaproteobacteria, and Bacteroidia. Community structure was related to water mass, salinity, and bloom stage, but not to depth. Clusters ranged from Gammaproteobacteria-dominant, Bacteroidia-poor assemblages in warm and cool Shelf Water (Division A) to Bacteroidia-rich assemblages spanning pre- to post-bloom conditions (Divisions B and C). In distance-based redundancy analysis, temperature, salinity, dissolved oxygen, nitrate, and fluorescence together explained 14.8 % of the variation in community composition, with salinity the strongest contributor. These results indicate that water-mass origin, physicochemical conditions, and bloom stage jointly shape bacterial communities, and that changes in temperature, salinity, and the magnitude and timing of blooms could substantially alter bacterial communities of this Pacific gateway to the Arctic.

## 1 Introduction

Microbes are the most abundant organisms in the global oceans, are essential in marine ecosystems, and include bacteria, archaea, algae, protists, fungi, and viruses. Marine organisms at higher trophic levels graze microbes and consume organic matter that is produced by microbes, and microbes contribute to marine nutrient cycling through the remineralization of organic matter into inorganic nutrients like nitrate and phosphate, which are then taken up by other organisms.

The Arctic Ocean is a habitat for many forms of charismatic fauna, but the fact that microbes are supporting their lives is often overlooked. To understand the role of microbes in the Arctic Ocean, studying their diversity, distribution, and function is necessary. The Arctic Ocean harbors unique marine ecosystems, and the loss of microbial diversity due to climate change is likely to cause shifts in microbial community structure, leading to the loss of unique microbial species and their influence on biogeochemical cycles and marine food webs in Arctic marine ecosystems (Comeau et al., 2011; Kraemer et al., 2024). Shifts in microbial community composition may also accompany the increased occurrence of harmful algal blooms in the region. In fact, massive harmful algal blooms (HABs) were recently observed in the Bering Strait (Fachon et al., 2025). HABs may also have disproportionate impacts on people in Indigenous communities who utilize the waters for subsistence livelihoods (Nolan, 2022). Therefore, studying microbial biodiversity is important because a loss of diversity could have unforeseen consequences, not only for the marine ecosystem itself but also for the communities that depend on it.

Microbial biogeography is the study of the distributions of microbes in time and space. Knowing the baseline microbial distributions at specific locations allows us to assess the effects of environmental change on microbial communities over time. For example, psychrophiles (cold-loving microbes) are abundant in the Arctic Ocean and can only survive in cold environments. If sea surface temperatures rise, the abundance of psychrophiles will likely decrease; monitoring their abundance will help us understand the effect of climate change on the Arctic microbial community. Also, since some psychrophiles in the Arctic Ocean are endemic to that region, their loss may act as a bioindicator of climate change and potentially provide forewarning of larger scale changes in ecosystem function. Microbial function includes the role of each microbe in biogeochemical cycling and marine food webs, including primary production, secondary production, and remineralization of inorganic species of carbon, nitrogen, phosphorus, and silicon. As mentioned above, microbes support marine organisms at higher trophic levels, so microbial compositional changes may influence the efficiency of food web functionality and lead to changes in the entire ecology, for example by changes in the phenology (timing) of algal blooms. In addition, studying microbial function helps us understand the mechanisms by which microbes evolved and adapted their metabolism and physiology to the constraints present in the cold, dark (or light), Arctic Ocean.

Finally, studying microbial function will benefit researchers in other fields as well, for example biogeo-chemical modelers observe that microbes are essential drivers of Arctic biogeochemical cycling, and that microbial compositional change can impact those cycles, which are ultimately linked to climate change. By studying microbial functionality in the Arctic Ocean, we will help determine the relative importance of microbes in the Arctic marine ecosystem, and whether there are unique aspects of Arctic microbial communities there compared to other oceans. Due to global warming, the Arctic region is warming four times faster than other regions (Serreze & Barry, 2011). Also, Comeau et al. (2011) and Kraemer et al. (2024) reported that microbial community composition in the Arctic Ocean changed dramatically after the huge loss of multiyear ice (MYI) in 2007 and 2012. Along with global warming, seasonal sea ice melt can also change microbial compositions in the Arctic Ocean. Freshwater input from sea ice melt during spring can influence microbial communities in the Arctic Ocean (Campbell et al., 2018) and microbial food webs (Kellogg et al., 2019) in Arctic estuaries. Those results imply that climate change may change the component of microbial community which leads to the loss of microbial diversity, unique microbes, and their unique metabolism.

Studying microbes in the Arctic Ocean will help us understand the Arctic marine ecosystem and assess the impact of climate change on the ecosystem. With metagenomic techniques, biogeography, metabolism, and diversity of Arctic marine microbes have been analyzed in the Arctic Ocean. However, compared with other oceans, we only have limited information about the Arctic marine microbial communities due to their relative inaccessibility. Therefore, in this study, we analyze the microbial community in the Bering and Chukchi Seas, which are at the entrance to the Arctic Ocean on the Pacific side and are known as areas with high productivity (Grebmeier, 2012; Grebmeier et al., 2006; Springer et al., 1996).

Here, using 16S rRNA gene amplicon sequencing of 181 samples collected at three depths (surface, middle, and bottom) across 62 stations during the Arctic Shelf Growth, Advection, Respiration and Deposition Rate Experiments (ASGARD) 2017 cruise, we aim to (i) characterize the composition and spatial structure of bacterial communities in the Bering and Chukchi Seas, (ii) examine how the diatom bloom influenced community composition, (iii) identify relationships between the communities and physicochemical parameters of water masses from which they derived, and (iv) assess how physicochemical changes associated with ongoing warming may affect these communities.

By linking community structure to bloom dynamics and to the physicochemical properties of water masses, this study provides a baseline for understanding how microbial communities in this region respond to environmental change and evaluates their potential as sentinels of that change.

## 2 Methods

### 2.1 Sample collection

Seawater sampling was conducted during the ASGARD project from June 9–28, 2017 aboard R/V Sikuliaq. The ASGARD 2017 cruise visited 62 “survey” stations (Figure 1), which were brief stops to collect water for nutrients, microbes, and conduct zooplankton tows. The ASGARD 2017 cruise visited 68 unique stations, many of which were associated with the Distributed Biological Observatory (DBO), including the DBO2 line in the northern Bering Sea and the DBO3 line in the southern Chukchi Sea. Samples were collected from surface, middle (pycnocline) and bottom depths between 2.5 m (surface) and 55 m (bottom), with modal depths per station at 5 m, 20 m, and 40 m. Seawater was sampled from Niskin bottles on a rosette with an attached Sea-Bird CTD (Sea-Bird Electronics Inc., Bellevue, WA, USA).

**Figure 1.**
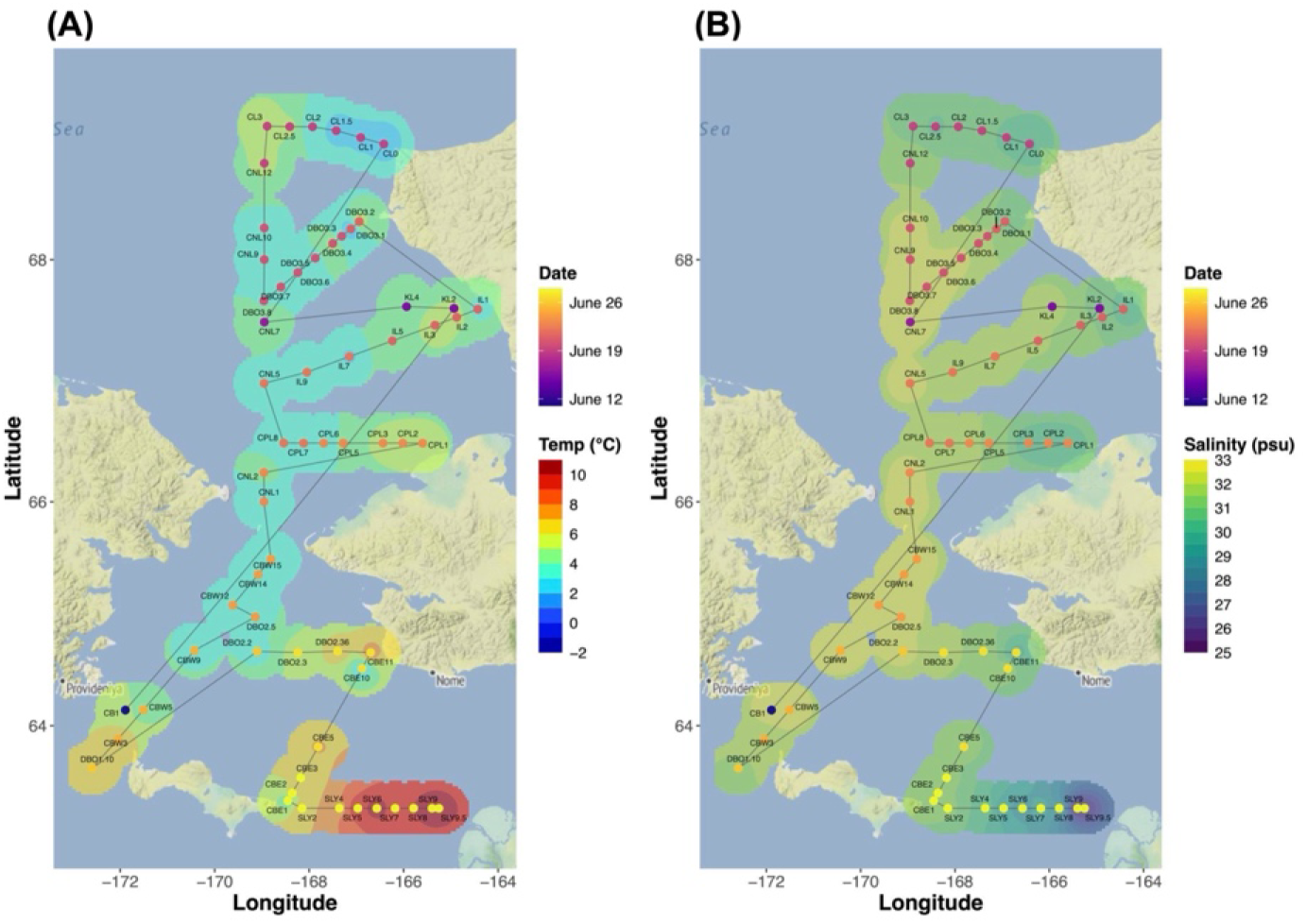
Survey Stations and Near-surface Hydrography During the ASGARD 2017 Cruise. Survey stations of the ASGARD 2017 cruise across the Bering and Chukchi Seas. Each station is colored by its sampling timing, is connected with the grey line in chronological order to indicate the cruise track, and is labeled by its station name based on pre-determined lines: SLY (St. Lawrence Island to Yukon River transect), CBE (Chirikov Basin East transect), CBW (Chirikov Basin West transect), DBO (Distributed Biological Observatory), CPL (Cowpatch Line transect), CNL (Convention Line transect), IL (Isilivik Lagoon transect), KL (Kivalina Line transect), and CL (Cape Lisburne transect). The filled background shows the near-surface (A) temperature and (B) salinity, interpolated from the station values by inverse-distance weighting and masked to within 30 km of a station.

Samples were collected in two phases, here labeled “survey” and “process” stations. The survey stations (62 total) were relatively short stops where only the CTD rosette and zooplankton nets were deployed; process stations (12 total) were full-day stops, including four CTD casts conducted for sensor profiling, nutrient analysis, zooplankton incubations, and more. Some stations were visited during both phases. Figure 1 shows the survey stations of ASGARD 2017 and the near-surface hydrography during the cruise obtained by the flow-through thermosalinograph. The average near-surface temperature was 4.9 °C (range 1.1–10.9 °C), and salinity 31.6 (range 25.0–32.9). Six stations (CBE1, CBW5, CL1, CL3, DBO3.3, and IL) were sampled in both the survey and process phases. All geographic maps were produced in R with the ggmap package (Kahle & Wickham, 2013). Basemap data are © Stadia Maps, © Stamen Design, © OpenMapTiles and © OpenStreetMap contributors.

### 2.2 Sample filtration

Seawater samples were transferred from Niskin bottles to 20 L cubitainers and stored at 4 °C until filtration within six hours. Using a peristaltic pump, 1–5 L of seawater per depth was filtered. Samples from the survey stations were directly filtered onto 0.2 µm-pore size Sterivex cartridge filters to collect microbes (Millipore Sigma, Burlington, MA, USA). Filters were stored at −20 °C until processing.

Nutrient samples were drawn with 60 mL syringes immediately after CTD recovery and filtered through 0.45 µm Nuclepore filters, and kept frozen below −20 °C until processed colorimetrically by autoanalyzer post-cruise (Gordon, Jennings and Krest 1993). Nitrate, phosphate and silicate concentrations were provided by S. Danielson (University of Alaska Fairbanks, unpublished data).

### 2.3 DNA extraction, amplification, and sequencing

DNA was extracted from filters using the DNeasy PowerWater kit (Qiagen, Hilden, Germany) following the manufacturer’s instructions. PCR-amplification of 16S rRNA genes was used for analysis of prokaryotic communities. The KAPA HiFi HotStart PCR Kit (Kapa Biosystems, Wilmington, MA, USA) was used for PCR-amplification of 16S rRNA genes using the following conditions: one cycle at 98 °C for 1 min, 26 cycles at 98 °C for 30 s, 55 °C for 30 s, 72 °C for 30 s, and one cycle at 72 °C for five minutes. Earth Microbiome Project primers (Thompson et al., 2017) were used to target the V4 hypervariable region of 16S SSU rRNA: forward primer 515F-revised (Parada et al., 2016) (5’-GTGYCAGCMGCCGCGGTAA-3’) and reverse primer 806RB-revised (Apprill et al., 2015) (5’-GGACTACNVGGGTWTCTAAT-3’). Amplified DNA was dual-indexed using unique adapters (Glenn et al., 2016) before TruSeq library preparation and sequencing on an Illumina MiSeq in the DNA Core Lab at the University of Alaska, Fairbanks (Fairbanks, USA).

### 2.4 Analysis of 16S rRNA gene sequencing

Sequences were demultiplexed and primers were removed with cutadapt v2.8 (Martin, 2014). The amplicon sequencing data was analyzed for amplicon sequence variants (ASVs) using DADA2 in R (Divisive Amplicon Denoising Algorithm 2; Callahan et al., 2016), performing quality control, error correction, merging, chimera checks and taxonomic classification. The SILVA database v132 (Quast et al., 2012) was used to classify ASVs.

In total, 3076 prokaryotic amplicon sequence variants (ASVs) were obtained from 16S rRNA gene and samples with fewer than 1000 reads were excluded. Of the 3076 ASVs, those detected with *≥* 2 reads in each dataset numbered 442 in the survey. Read counts were converted to relative abundances within each sample and ASVs were retained only if their maximum relative abundance across samples exceeded 0.001 (0.1 %) and they occurred in two or more samples. This filtering yielded 258 ASVs from the 181 survey samples.

The analyses described in the following sections were performed in R (version 4.5.2). All R scripts and outputs are publicly available at https://github.com/ReiSHIBUE/ASGARD2017_analysis.

### 2.5 Hierarchical Clustering

For beta diversity analysis, Bray-Curtis dissimilarities were calculated from the fourth-root-transformed relative abundance matrix (258 ASVs in 181 survey samples) and hierarchically clustered using Ward’s minimum variance method. Dissimilarities were calculated with the vegan package in R (Oksanen et al., 2024), and the clustering was plotted with the R packages vegan, fastcluster (Müllner, 2013), and gplots (Warnes et al., 2009). Clusters were defined by cutting the dendrogram, and labels were assigned hierarchically following the dendrogram topology: each top-level branch was given a division-level name, and within each division the subsequent splits defined nested subclusters (e.g., division-level *→* first split *→* second split). For the survey dataset, the resulting dendrogram was cut at k = 10 and the groups were separated into three divisions (A, B, C), yielding a final set of 11 clusters (A1, A2, B1, B2a, B2b, C1a, C1b1, C1b2, C2a, C2b1, and C2b2) following manual separation of C2b into C2b1 and C2b2.

For the further separation in Division C, a top-down tree walk was conducted and the dendrogram was interpreted against an environmental ordination. The nine environmental variables (temperature, salinity, DO, NO_3_, PO_4_, silicate, NH_4_, fluorescence and depth) were standardized and ordinated by principal component analysis (PCA), and the first nine components (PC1–PC9) were retained and interpreted from their loadings. Division C clusters were subdivided, and at each split of the dendrogram, the two descendant subclusters were compared along each PC separately: for every PC, a one-dimensional Euclidean distance was computed from the sample PC scores and tested by pairwise permutational multivariate analysis of variance (PERMANOVA) with 999 permutations, with permutational analysis of multivariate dispersions (PERMDISP) used to check homogeneity of dispersion. Significance of PERMDISP was assessed by Tukey’s honestly significant difference (HSD) test. Within each node, the p-values across the nine PCs were BH-adjusted, and the principal component(s) showing the most significant separation were regarded as the environmental axis driving that split.

Bubble plots were generated to visualize relative abundances of ASVs belonging to three major bacterial classes (Alphaproteobacteria, Gammaproteobacteria, and Bacteroidia). For each sample, ASV relative abundances were summed to the genus level and averaged across the samples within each cluster, giving a mean relative abundance (%) per genus per cluster.

### 2.6 Water Mass Characterization

A temperature–salinity (T-S) diagram was constructed to characterize the water masses sampled and to visualize their relationship with the microbial community clusters. Potential density isopycnals were overlaid for reference, and water mass boundaries followed the classification of Danielson et al. (2020), reproduced in Table 1. All plots were produced with ggplot2 in R (Wickham, 2016). Seawater density isolines were computed from the EOS-80 equation of state. The correlation between each cluster and water mass was assessed by Fisher’s exact test (9,999 Monte Carlo replicates), while the significance of correlations between each cluster and depth was tested with a Kruskal–Wallis test.

**Table 1.** Water Mass Definitions in the Northern Bering and Chukchi Seas, Modified From Danielson et al. (2020)

| Water mass | Abbreviation | Temperature range (°C) | Salinity range |
|---|---|---|---|
| Anadyr Water | AnW | 0 < T < 3 | 32.5 < S < 33.8 |
| Ice Melt Water & cool Coastal Water | IMW, cCW | −2 < T < 3 | 22 < S < 30.8 |
| cool Shelf Water | cSW | 0 < T < 3 | 30.8 < S < 32.5 |
| warm Coastal Water | wCW | 3 < T < 14 | 18 < S < 30.8 |
| warm Shelf Water | wSW | 3 < T < 14 | 30.8 < S < 33.4 |
| Modified Winter Water | MWW | −1 < T < 0 | 30.8 < S < 33.8 |
| Winter Water | WW | −2 < T < −1 | 30.8 < S < 35 |

*Note.* Atlantic Water and Bering Basin Water (AtlW and BBW) are not included in this table because no samples in this study fell within their temperature–salinity ranges. Temperature (T) is in °C and salinity (S) is unitless (PSU).

Samples were assigned to the Bering or Chukchi by latitude (south or north of the Bering Strait at 65.7 °N, respectively), and potential density was calculated from temperature and salinity using the EOS-80 equation of state at surface pressure.

### 2.7 Environmental Characterization of Sample Clusters

Boxplots were generated for each measured environmental variable (temperature, salinity, dissolved oxygen, nitrate, phosphate, ammonium, chlorophyll fluorescence, chlorophyll a, sampling depth, and latitude) to explore how environmental conditions varied among microbial community clusters. Each cluster was visualized with median, interquartile range, and overlaid individual sample points using ggplot2 in R.

To classify each microbial community cluster by its bloom stage, principal component analysis (PCA) was performed on nine standardized environmental variables (with log(1+x)-transformation of right-skewed nutrient and fluorescence variables). PC1 was observed to capture the dominant gradient between pre- and post-bloom conditions, and was rescaled to the [0, 1] interval and interpreted as a “bloom index” (0 = pre-bloom, 1 = post-bloom). The index is derived from environmental data alone and uses no microbial information, so the assignment of bloom stages to clusters is independent of the clustering itself. The distribution of the bloom index across clusters was then used to assign a representative bloom stage to each cluster.

Parameter selection was conducted by calculating variance inflation factors (VIF) between each environmental parameter using the R package car (Fox & Weisberg, 2019) and GGally (Schloerke et al., 2024). Temperature (°C), salinity, depth (m), DO (µmol/kg), NO_3_ (µM) and FlECO-AFL (chlorophyll fluorescence, mg m^−3^) were selected for further analysis. A distance-based redundancy analysis (dbRDA) was performed, and ordinations were used to conduct ANOVA between environmental parameters and microbial ASVs using the vegan package. dbRDA plots were generated using ggplot2. Bray–Curtis distances were calculated from the fourth-root-transformed ASV relative abundance matrices. Five parameters were used for this analysis (temperature, salinity, DO, NO_3_, and FlECO-AFL) and only samples with complete environmental data were analyzed. The significance of the overall model, the marginal contribution of each variable, and each constrained axis were tested by permutational ANOVA with 999 permutations and the constrained variance was reported as the adjusted R². Ordinations were visualized with ggplot2, with the environmental variables displayed using stats::biplot.

ASV richness, defined as the number of ASVs present (relative abundance > 0) in the filtered clustering set, was compared across each depth using Kruskal–Wallis rank-sum tests. Tests were performed overall and separately within each community grouping, followed by Bonferroni correction for the number of within-group tests.

## 3 Results

### 3.1 Hierarchical Clustering

In total, 181 samples were assigned to three divisions (A, B, and C) and those were further categorized into 11 clusters based on the dendrogram (Figure 2): Cluster A1 (red; n = 9), Cluster A2 (orange; n = 15), Cluster B1 (dark green; n = 17), Cluster B2a (light green; n = 17), Cluster B2b (yellow-green; n = 26), Cluster C1a (dark blue; n = 14), Cluster C1b1 (purple; n = 11), Cluster C1b2 (brown; n = 19), Cluster C2a (light blue; n = 12), Cluster C2b1 (lavender; n = 12), and Cluster C2b2 (gray; n = 29). In addition, 258 filtered-ASVs were grouped into six Assemblages: Assemblage 1 (black; # of ASVs = 24), Assemblage 2 (pink; # of ASVs = 27), Assemblage 3 (yellow; # of ASVs = 25), Assemblage 4 (light blue; # of ASVs = 30), Assemblage 5 (dark blue; # of ASVs = 52), and Assemblage 6 (light red; # of ASVs = 100).

**Figure 2.**
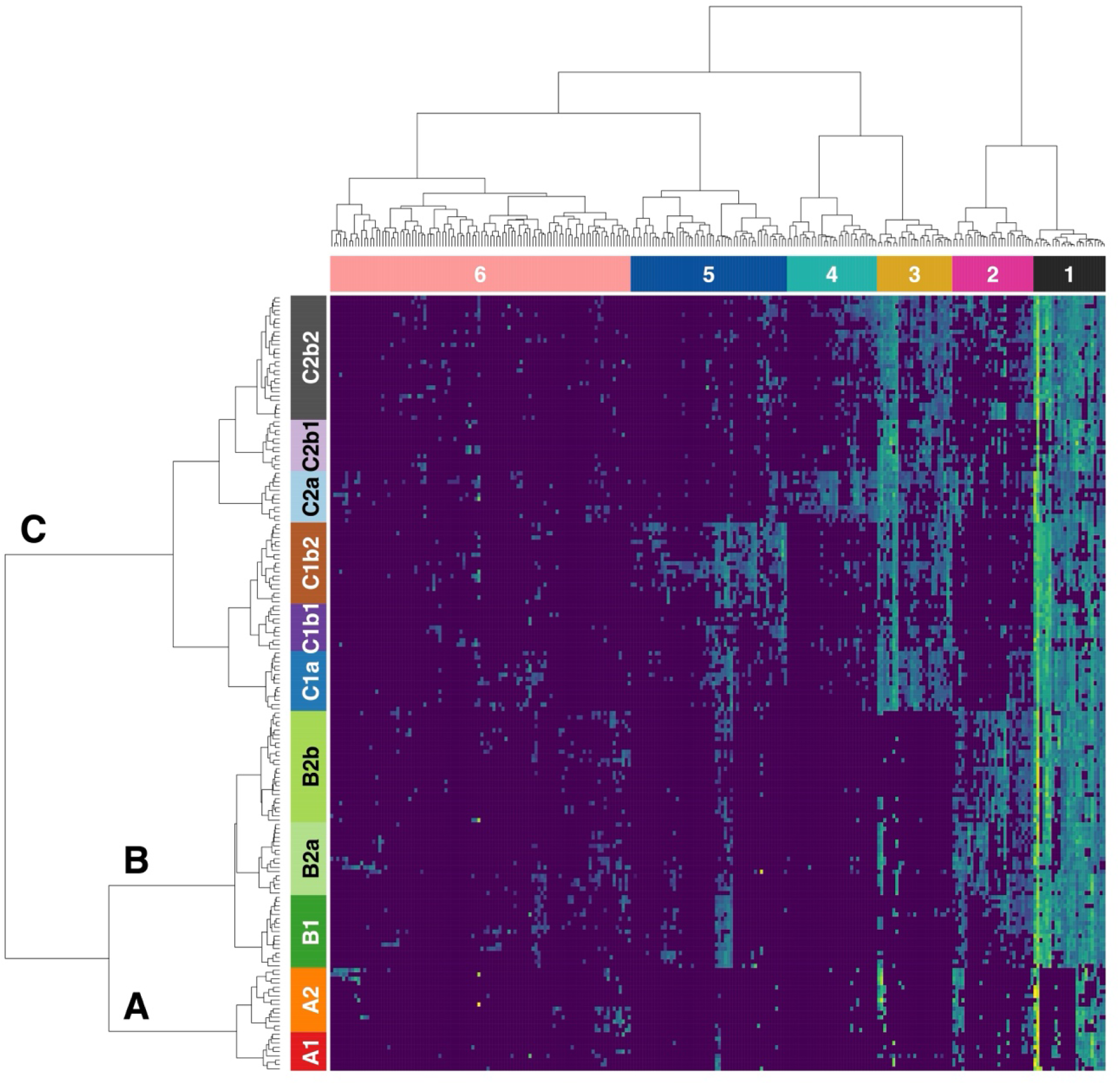
Heatmap of ASVs in the Survey Dataset. Hierarchical clustering was conducted based on relative abundance of 258 ASVs (columns), present in at least two of 181 samples (rows) with maximum relative abundances greater than or equal to 0.1 %. Samples were assigned to three primary divisions (A, B, and C) in 11 clusters based on the dendrogram (Cluster A1 to C2b2). ASVs were assigned to one of six assemblages (columns) based on the dendrogram.

Likewise, ASVs were assigned to one of six assemblages based on clustering (Figure 2). Mean relative abundance of each ASV was calculated, revealing that uncultured Nitrincolaceae, *Sulfitobacter*, SAR11 clade Ia, *Polaribacter*, *Ulvibacter*, and OM182 clade were highly abundant throughout all sample clusters and were mostly assigned to Assemblage 1. Therefore, we categorized them as the cosmopolitan ASVs.

Clusters A1 and A2 were distinguished by their low diversity and lack of numerous ASVs from Assemblage 1 assigned to the Bacteroidetes, including *Polaribacter*, *Ulvibacter* (two ASVs), uncultured Cryomorphaceae, uncultured Flavobacteriaceae, *Tenacibaculum*, NS11-12 marine group and *Fluviicola*. A2 and B2a were differentiated from A1 and B2b respectively by showing high abundance of two SAR86 clade ASVs.

Division B clusters (B1, B2a, and B2b) were defined by ASVs in Assemblage 2, which were mostly dominated by Bacteroidetes such as *Polaribacter* (three ASVs), NS4 marine group (two ASVs), *Lutibacter* (two ASVs), *Lutimonas*, and uncultured Cryomorphaceae (two ASVs), representing particle-attached copiotrophs associated with phytoplankton blooms and Gammaproteobacteria categorized as OMG (Oligotrophic Marine Group) such as SAR92 (four ASVs), OM182 clade (two ASVs), and OM60/NOR5 clade.

Division C clusters (C1a, C1b1, C1b2, C2a, C2b1 and C2b2) were characterized by ASVs in Assemblage 3, which included numerous Bacteroidetes ASVs (predominantly NS marine groups ASVs), OMG group (SAR86 clade and OM60/NOR5), methylotrophs (OM43 clade, *Methylotenera*, and Methylophagaceae) and other heterotrophic marine bacteria. C1 groups (C1a, C1b1, and C1b2) were distinct from C2 groups (C2a, C2b1, and C2b2) by taxonomically diverse ASVs in Assemblage 5, including *Colwellia*, Marine Group II archaea, and other ASVs with low bootstrap values, suggesting unreliability of taxonomic annotations in this column cluster. ASVs in Assemblage 5 also divided other clusters in Division C. For instance, Division B and C1 group shared some ASVs including sulfur-oxidizing bacteria ASVs (two Thiotrichales ASVs) and NS4 and NS9 marine groups ASVs. Also, C1b1 and C1b2 were separated by ASVs, comprised of various bacteria including Flavobacteriales ASVs (NS marine groups, uncultured Cryomorphaceae, and *Fluviicola*), chemolithoautotrophic bacteria (SUP05 cluster and SAR324), and oligotrophic clades (SAR86). C2b1 and C2b2 were also divided by ASVs such as sediment associated bacteria (*Illumatobacter* and Urania-1B-19 group), *Colwellia*, SAR11 clade II, NS7 marine group, and *Alteromonas*. C2a and C2b1/C2b2 were divided by taxonomically distinct ASVs in Assemblage 4, including Flavobacteriales (NS5/7 marine groups and Cryomorphaceae), oligotrophs (SAR11 clade II/III/IV and AEGEAN-169), methylotroph (OM43 clade), chemolithotrophs (Nitrosomonadaceae, Thiotrichaceae, and Chromatiales), and Cyanobacteria (*Atelocyanobacterium* (UCYN-A)).

ASVs in each assemblage showed different patterns (Figure 3). ASVs in Assemblage 1 were highly abundant throughout the Bering and Chukchi Seas and its mean relative abundance was 70.3 %. Those ASVs exhibited a cosmopolitan-like distributional pattern and globally dominant taxa such as Nitrincolaceae and SAR11 clade Ia (Giovannoni, 2017; Thiele et al., 2022; Wietz et al., 2021) were highly dominant in this Assemblage. This constant distributional pattern in both seas may be attributed to the fact that both seas share the same Pacific-origin water. One study reported that microbial communities above 44°N showed similar community structures (Milke et al., 2022). ASVs in Assemblage 2 and 5 showed different patterns. For example, Assemblage 5 ASVs showed positive correlation with NO_3_ on dbRDA analysis (F = 21.4, p = 0.001, permutations = 999, adjusted R² = 0.37) while Assemblage 2 ASVs showed negative correlations with it (F = 19.6, p = 0.001, permutations = 999, adjusted R² = 0.35). This difference can be attributed to whether abundant ASVs were found in the area where Anadyr Current flows. The origin of AnW is nutrient-rich water from deep (Coachman et al., 1975) and ASVs in Assemblage 5 were more abundant in that area. Also, *Nitrosopumilus*, which is a well-known ammonia oxidizer in deep water (Könneke et al., 2005), was relatively more abundant in this Assemblage. On the other hand, ASVs in Assemblage 2 were more abundant in the east which is influenced by Alaskan Coastal Current and Bering Shelf Current. Therefore, both assemblages showed different correlation with NO_3_. ASVs in Assemblage 4 were more abundant around the Alaskan coast. Those ASVs were abundant in Cluster C2a, which showed lower salinity than other clusters (Figure 7). Also, ASVs in Assemblage 3 showed cosmopolitan-like distribution and did not show specific correlations with environmental variables on dbRDA. However, most of them were collected with 0.2 µm filter, while ASVs in Assemblage 6 were mainly collected with 20 µm filter (Figure 4). This result suggested that Assemblage 3 is largely composed of free-living microbes, while Assemblage 6 is mostly composed of particle-attached microbes. ASVs in Assemblage 3 separated Division C clusters and were mainly composed of Bacteroidia, OMG group, and methylotrophs. On the other hand, 100 ASVs were assigned to Assemblage 6, but they did not show specific abundance pattern on heatmap (Figure 2). This result implies that Assemblage 6 is a patchy group of ASVs from the rare biosphere and cannot be represented by any single dominant ASV.

**Figure 3.**
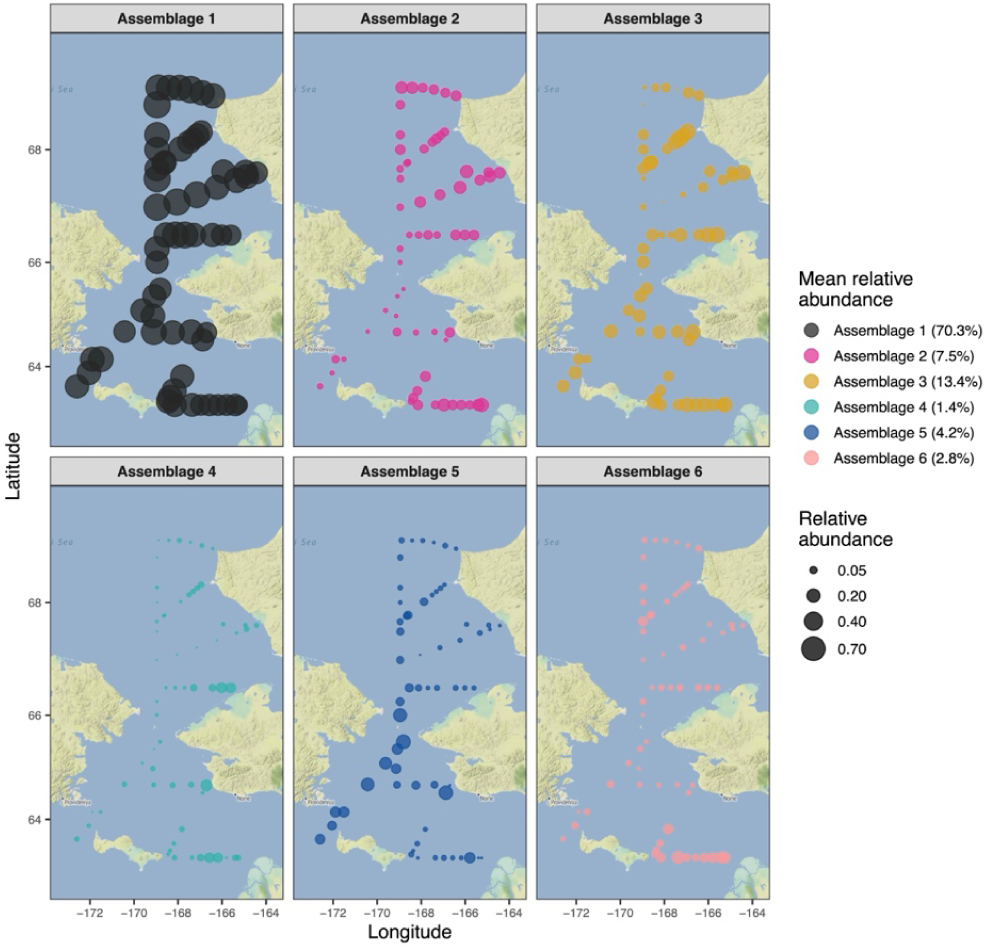
Sampling Sites of ASVs in Six Assemblages in the Survey Dataset. Each Assemblage is assigned to different colors. The dots correspond to mean relative abundance of ASVs in six assemblages. Mean relative abundance of ASVs in each Assemblage is shown as legend.

**Figure 4.**
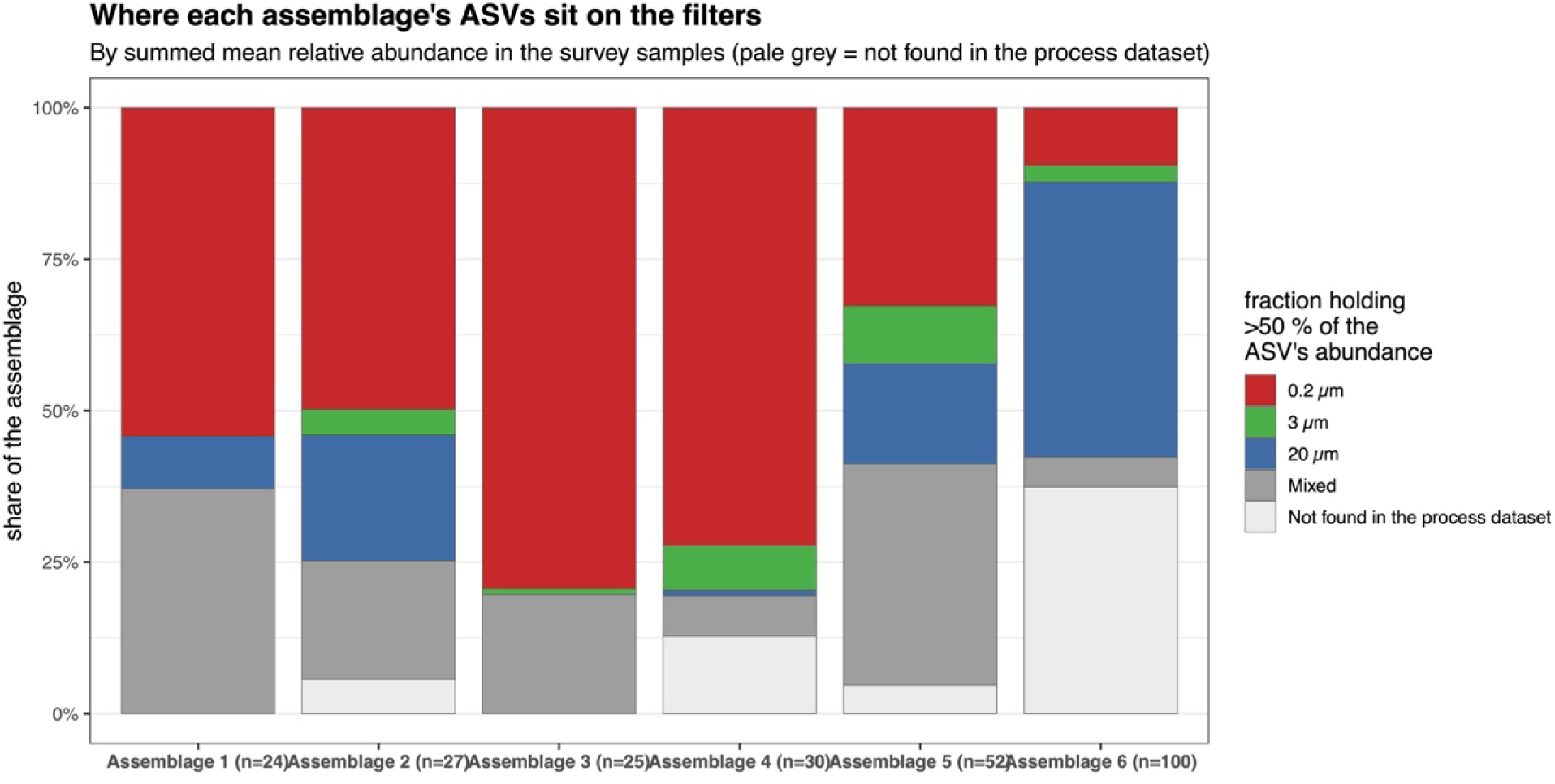
Relative Abundance of Six Assemblages ASVs With Each Filter in the Survey Dataset. ASVs whose relative abundance was greater than 50 % were assigned to each filter size. Each filter is represented as different colors (red: 0.2 µm, green: 3 µm, blue: 20 µm). Gray color means that ASVs whose abundance is spread across the size fractions, with no single filter holding more than 50 % of the ASV’s abundance. White color means that ASVs are absent from the process dataset and thus, it cannot be assigned to any size fractions.

Overall PERMANOVA indicated significant differences in community composition among the 11 clusters (R² = 0.587, F = 24.13, p < 0.001), while PERMDISP also revealed significant heterogeneity in within-cluster dispersion (F = 7.65, p < 0.001). Since PERMANOVA is sensitive to both differences in group centroids and within-group dispersion, the co-significance of PERMDISP suggests that the PERMANOVA result may be partly influenced by unequal dispersions, particularly for pairs involving Cluster A2.

Therefore, pairwise PERMANOVA and PERMDISP were conducted to determine which specific cluster pairs differed in composition and which clusters were responsible for the dispersion heterogeneity. Pairwise PERMANOVA revealed that all cluster combinations had significantly distinct microbial communities (average R^2^ = 0.4182, BH-adjusted p < 0.001). On the other hand, pairwise PERMDISP was non-significant in 44 of the 45 pairwise comparisons not involving A2 (the exception was C1a vs A1, Tukey’s HSD p = 0.037), indicating that PERMANOVA significance for these pairs can largely be attributed to true differences in community composition rather than dispersion. Cluster A2 was the main source of this heterogeneity, exhibiting the highest mean distance to centroid (0.336) and significantly greater dispersion than other clusters except for Cluster A1 (Tukey’s HSD, p < 0.05), suggesting that A2 encompasses a broader range of microbial community compositions compared to other clusters. However, it should also be noted that A2 had the lowest median sequencing depth (5,470 reads) among the 11 clusters, which may have inflated Bray-Curtis dissimilarities and contributed to the elevated dispersion.

### 3.2 Geographic and Hydrographic Distribution of Sample Clusters

The geographic distribution of the 11 clusters revealed distinct spatial patterns (Figure 5). Division A clusters (A1 and A2) were largely confined to the southern portion of the study area. Especially, Cluster A2 was restricted to the stations in St. Lawrence to Yukon River Transect (SLY). Division B clusters were broadly distributed across the Bering and Chukchi regions and Cluster B2b spanned the widest geographic range from Cape Lisburne Transect (CL) to Chirikov Basin East Transect (CBE) (16 stations). Within Division C, Cluster C1a was the most geographically restricted cluster, occurring exclusively at DBO3 stations (four stations) near the northern Chukchi Sea. In contrast, Cluster C2b2 had the broadest distribution (21 stations), which like Cluster B2b spanned the entire study area from south to north. Clusters C1b1 and C1b2 were mainly concentrated in Chirikov Basin West (CBW) and DBO1–2 stations, while Clusters C2a and C2b1 were mostly found in CBE and Cowpatch Line Transect (CPL). No significant association was detected between microbial communities in each cluster and depth types (Fisher’s exact test, p = 0.94), indicating that the 11 clusters were not driven by sampling depth.

**Figure 5.**
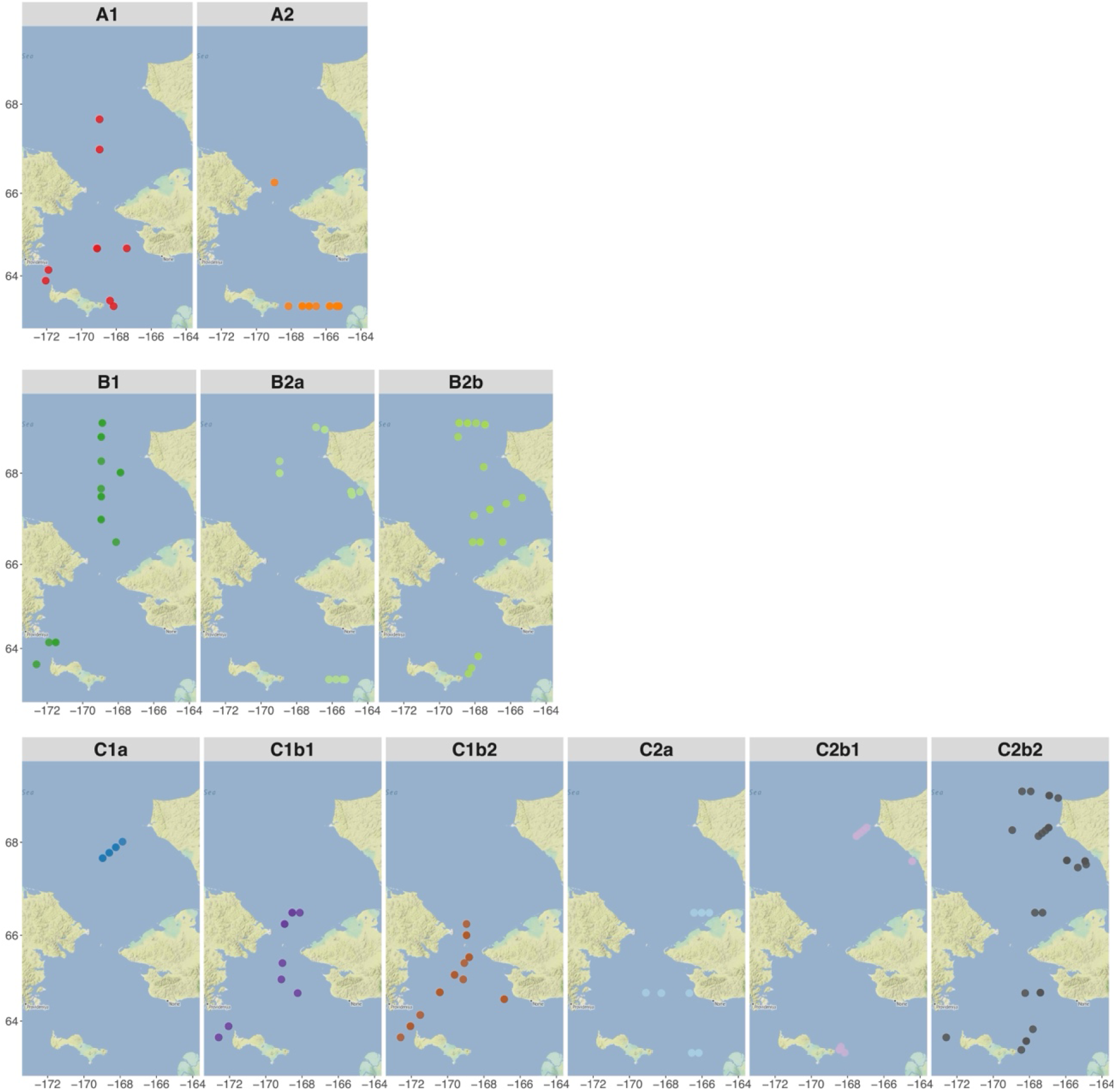
Maps of Sampling Stations for Each of the 11 Sample Clusters. Sampling stations of the eleven clusters are listed. Cluster A1 (red; n = 9), Cluster A2 (orange; n = 15), Cluster B1 (dark green; n = 17), Cluster B2a (light green; n = 17), Cluster B2b (yellow-green; n = 26), Cluster C1a (dark blue; n = 14), Cluster C1b1 (purple; n = 11), Cluster C1b2 (brown; n = 19), Cluster C2a (light blue; n = 12), Cluster C2b1 (lavender; n = 12), and Cluster C2b2 (gray; n = 29).

To examine whether the spatial distribution of the 11 clusters corresponded to distinct water masses, a T-S diagram of all survey samples was plotted (Figure 6), and each sample cluster overlaid on the water mass definitions from Danielson et al. (2020). The proportion of samples classified into each water mass was calculated for each cluster from the contingency table. Table 2 shows the water masses from which samples in each cluster were primarily and secondarily collected. The statistical significance of the association between clusters and water masses was highly significant (Fisher’s exact tests with simulated p-values, 9,999 Monte Carlo replicates, p < 0.001).

**Figure 6.**
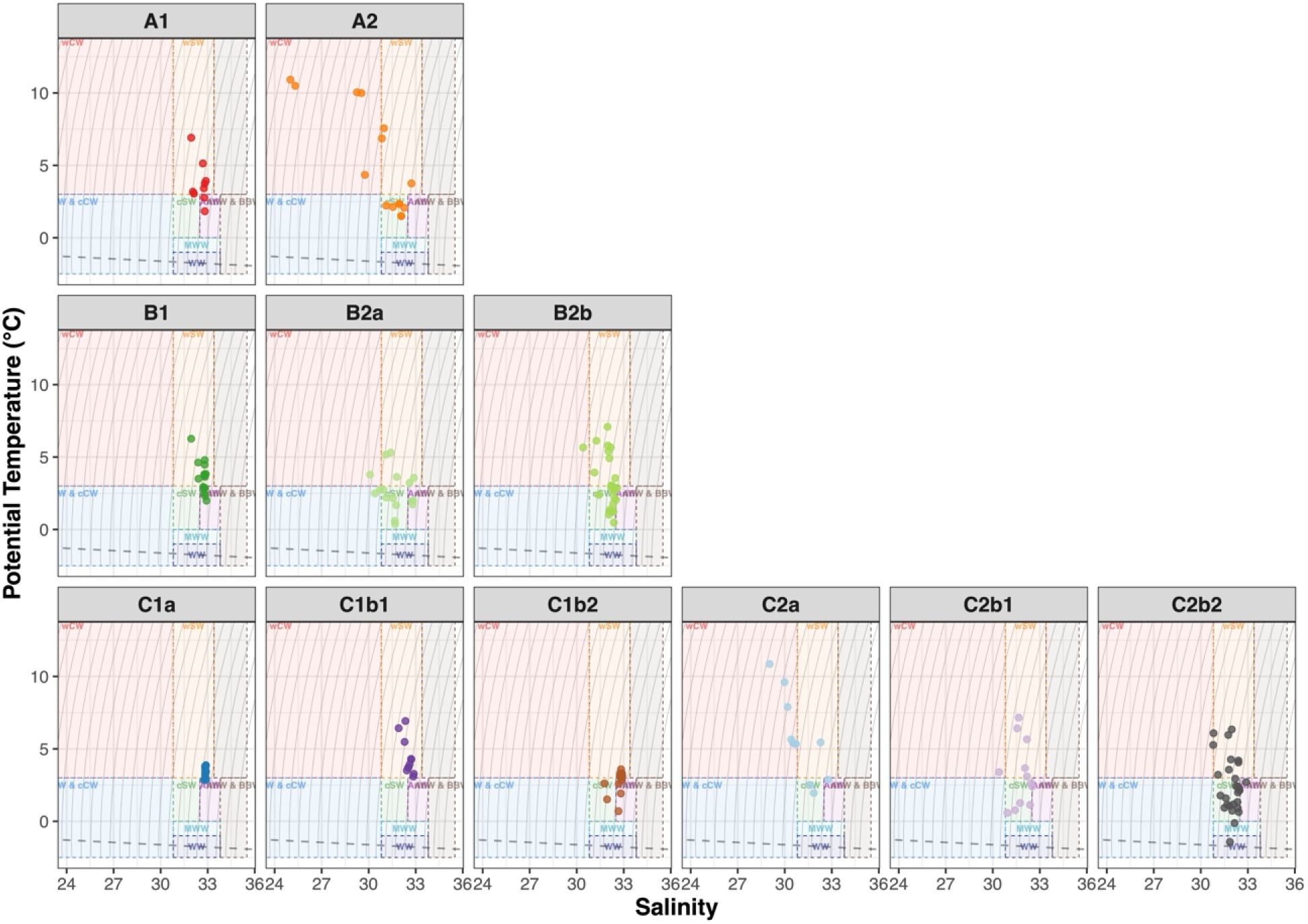
Water Mass Characteristics of the 11 Sample Clusters. Water mass definitions were adapted from Danielson et al. (2020). T-S diagram with water mass boundaries was separated by the 11 clusters. Dashed rectangles indicate the T-S ranges of each water mass: warm Coastal Water (wCW), cool Coastal Water (cCW), Ice Melt Water (IMW), warm Shelf Water (wSW), cool Shelf Water (cSW), Anadyr Water (AnW), Modified Winter Water (MWW), Winter Water (WW), Atlantic Water (AtlW), Bering Basin Water (BBW). Gray contours represent isopycnals (lines of equal density, *σ_θ_*, at 0.5 kg/m³ intervals). The dashed black line indicates the freezing point.

**Table 2.** The Proportion of Samples in the 11 Clusters, Classified Into Each Water Mass.

| Cluster | Primary water mass | Secondary water mass |
|---|---|---|
| A1 | wSW (78 %) | AnW (22 %) |
| A2 | cSW (47 %) | wCW (33 %) |
| B1 | wSW (53 %) | AnW (47 %) |
| B2a | cSW (35 %) | wSW (29 %) |
| B2b | cSW (54 %) | wSW (35 %) |
| C1a | wSW (79 %) | AnW (21 %) |
| C1b1 | wSW (100 %) | — |
| C1b2 | wSW (63 %) | AnW (26 %) |
| C2a | wCW (67 %) | cSW (17 %) |
| C2b1 | cSW / wSW (42 %) | — |
| C2b2 | cSW (59 %) | wSW (28 %) |
| Fisher’s exact test (Monte Carlo) |  |  |
| Replicates | *p* | Significance |
| B = 9,999 | 0.0001 | *** |

*Note.* The primary water mass was defined as the water mass with the highest proportion of samples, and the secondary water mass as the one with the second highest proportion. Water-mass abbreviations are defined in Table 1. An em dash (—) indicates that no second water mass was present. \*\*\* *p* < 0.001.

Most samples in each cluster were found from warm Shelf Water (wSW, n = 81, 44.8 %), cool Shelf Water (cSW, n = 53, 29.3 %), and Anadyr Water (AnW, n = 26, 14.4 %). Clusters A1, C1a, and C1b2 were primarily found from wSW and secondarily from AnW. Cluster B1 was mainly sampled from the same water masses, but samples were collected in roughly equal proportions from wSW and AnW. Many samples in Clusters A2, B2a, B2b, C2b1, and C2b2 were found from shelf waters (cSW and wSW), while Cluster C2a was the only cluster which was mainly sampled from warm Coastal Water (wCW). Cluster A2 occupied the broadest T-S range (salinity 25.0 – 32.7, temperature 1.5 – 10.9 °C), spanning cSW and wCW (Table 2), which may explain its elevated dispersion in pairwise PERMDISP.

### 3.3 Physicochemical Characterization of Sample Clusters

Physicochemical characteristics of the 11 clusters were shown in Figure 7. Clusters within Division A were sampled from relatively shallow depths (median: A1, 4.2 m; A2, 10.3 m), and A2 showed relatively low salinity (median: 31.1), reflecting the influence of wCW. Nutrient concentrations in Division A were generally low, with A2 showing near-complete depletion of NO_3_ (median: 0.1 µM). Division B clusters exhibited contrasting nutrient conditions. Cluster B1 had high concentrations of NO_3_ (median: 7.7 µM), suggesting active organic matter remineralization by heterotrophic bacteria. In contrast, B2a and B2b were nutrient-depleted (NO_3_ median: 0.15 and 0.4 µM, respectively), indicating post-bloom conditions.

**Figure 7.**
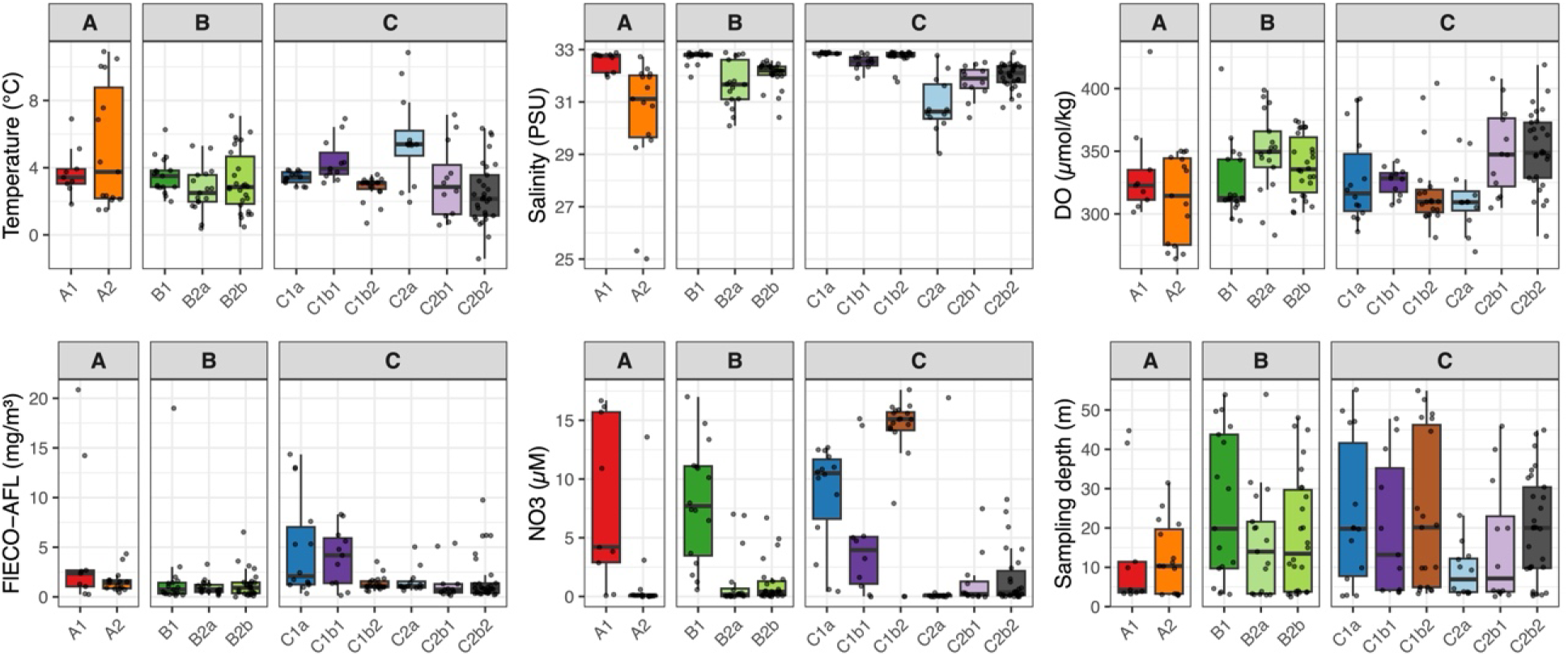
Boxplots of Environmental Variables for the 11 Sample Clusters, Grouped by Primary Divisions (A, B, and C). Each panel shows one of six variables: temperature (°C), salinity (PSU), DO (µmol/kg), FlECO-AFL (chlorophyll fluorescence, mg/m^3^), NO_3_ (µM), and sampling depth (m). Individual samples were overlaid as jittered points. Boxes represent the interquartile range (IQR), with the horizontal line indicating the median. Whiskers extend to the most extreme values within 1.5 × IQR.

Division C displayed the greatest environmental heterogeneity. Clusters C1b2 and C1a had the highest nutrient concentrations (NO_3_ median: 15.1 and 10.5 µM, respectively), consistent with pre-bloom conditions where nutrients had not yet been consumed by phytoplankton. In contrast, Cluster C1b1 exhibited the highest FlECO-AFL (median: 4.2 mg/m³) values among all clusters, suggesting active phytoplankton bloom conditions. C2 subclusters (C2a, C2b1, and C2b2) were uniformly nutrient-depleted (NO_3_ median: 0.1 – 0.4 µM), which is characteristic of post-bloom environments. Among these, C2a had the highest temperature (median: 5.4 °C) and the lowest salinity (median: 30.6), reflecting wCW influence (Table 2). On the other hand, C2b2 had the lowest temperature (median: 2.1 °C), consistent with its association with cSW (Table 2). Each cluster contains samples from all three depths. Therefore, part of the spread shown here comes from vertical differences, not from differences between communities. This applies most to salinity and NO_3_, which both change with depth. The clusters did not differ in how their samples were distributed among depths (Fisher’s exact test, p = 0.94), so comparisons between clusters should not be affected. Even so, showing each variable by cluster and by depth together would give a more complete picture, and we leave this for future work.

Nine environmental variables with near-complete data coverage (*≤* 3.9 % missing values; 174 out of 181 samples) were used for PCA: temperature, salinity, DO, NO_3_, PO_4_, Sil, NH_4_, FlECO-AFL, and sampling depth. Variables with higher proportions of missing data (e.g., chlorophyll a, 10.5 %; POC/PON, 35.4 %) were not included to maximize the number of samples retained in the analysis. PC1 (48.1 % of total variance explained) was interpreted as a “bloom index”, as it was primarily driven by positive loadings of nutrient concentrations (NO_3_, PO_4_, NH_4_, and Sil), salinity, and depth, reflecting the progression of phytoplankton bloom. High PC1 values indicated pre-bloom (nutrient-replete) conditions, while low PC1 values indicated post-bloom (nutrient-depleted) conditions.

The bloom index, which is derived from environmental variables only, revealed distinct bloom stages within each division (Figure 8). Cluster A1 (median: 0.39) represented pre-bloom conditions since NO_3_ concentration was higher than Cluster A2 (median: 0.59), which showed post-bloom conditions (Figure 7). Cluster B1 (median: 0.23) was the most pre-bloom cluster, with high residual nutrient concentrations, while Cluster B2a (median: 0.59) and Cluster B2b (median: 0.49) were in post-bloom stages. Division C exhibited the widest range of bloom stages: Cluster C1b2 (median: 0.16) and Cluster C1a (median: 0.16) were the most pre-bloom clusters among all 11 clusters, while Cluster C2a (median: 0.67), Cluster C2b1 (median: 0.66), and Cluster C2b2 (median: 0.49) were in post-bloom stages. Cluster C1b1 (median: 0.38) was categorized into the pre-bloom cluster, but it can also be represented as a during-bloom cluster since it showed the highest median of chlorophyll a concentration (8.03 µg/l) and higher median of NO_3_ (3.95 µM) than other Division C post-bloom clusters (C2a, C2b1, and C2b2).

**Figure 8.**
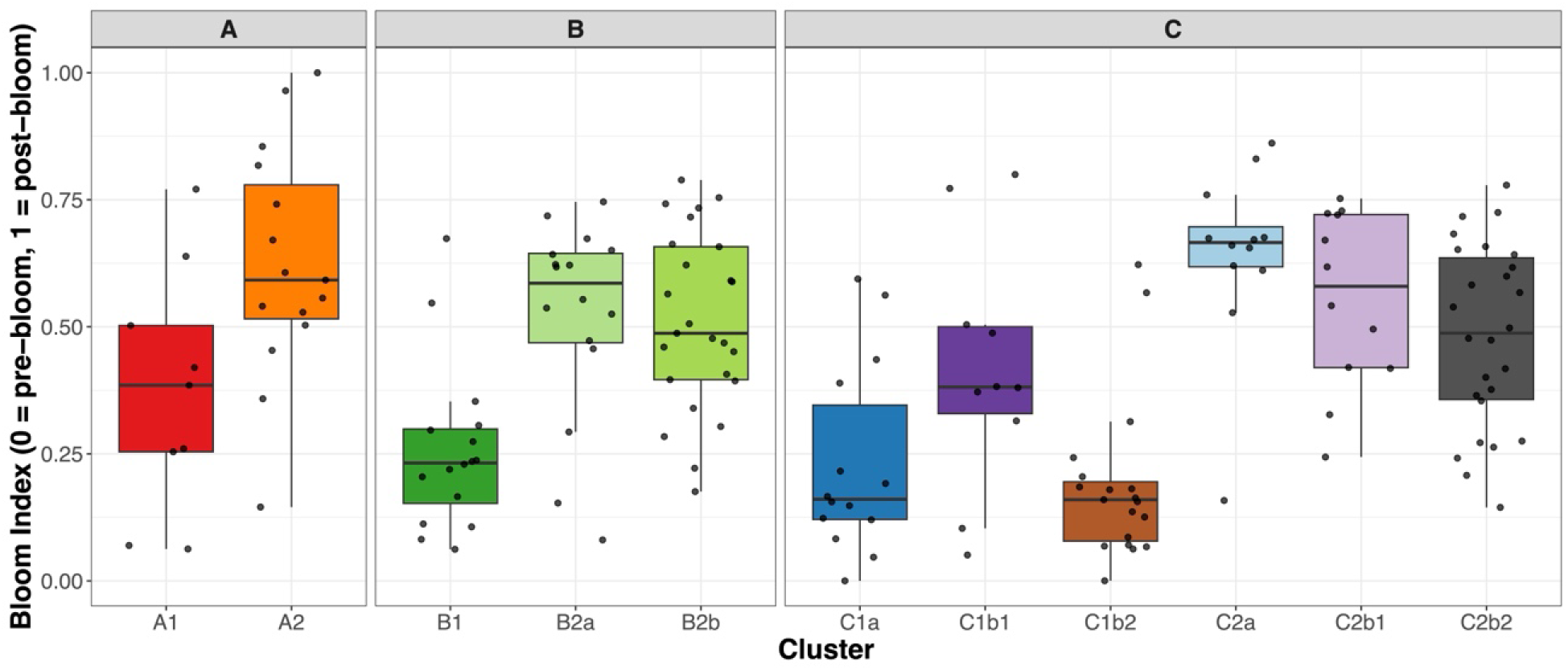
Bloom Index (Rescaled PC1) for the 11 Sample Clusters, Grouped by Primary Divisions (A, B, and C). The bloom index was derived from PC1 of a PCA on nine environmental variables, rescaled from 0 (pre-bloom, nutrient-replete) to 1 (post-bloom, nutrient-depleted). Boxes represent the interquartile range with the median shown as a horizontal line. Individual samples are overlaid as jittered points.

Since Division C (n = 97) was the most environmentally heterogeneous division, encompassing the full range of bloom stages from pre-bloom (C1a, C1b1, C1b2) to post-bloom (C2a, C2b1, and C2b2) and spanning six water masses (wSW, cSW, AnW, wCW, MWW, and WW), pairwise PERMANOVA was conducted on each PC axis separately, with p-values corrected for multiple testing within each node (Benjamini-Hochberg correction across 9 PC axes; Figure 9). The primary split between C1 and C2 was best explained by PC1 (R² = 0.34, p = 0.009), representing the nutrient gradient (the bloom index). The separation between C1a and C1b was mainly associated with PC6 (R² = 0.24, p = 0.027), reflecting the NH_4_–salinity–depth gradient, although PC6 explained only 3.2 % of the total environmental variance. The differentiation of C1b1 from C1b2 was most strongly associated with PC5 (R² = 0.70, p = 0.023), which represents a silicate–salinity–temperature gradient, though PC5 accounted for only 5.5 % of the total variance. Although the splits of C1a vs C1b and C1b1 vs C1b2 were explained by PC axes with relatively low explained variance (PC6, 3.2 %; PC5, 5.5 %), it suggested that the finer-scale subdivisions within Division C were driven by relatively minor environmental gradients not captured by the dominant PC axes. The split of C2a from C2b was driven by PC2 (R² = 0.32, p = 0.014), the DO–temperature gradient. No PC axis significantly differentiated C2b1 from C2b2 (all BH-adjusted p > 0.88), suggesting that this split was driven by differences in microbial community composition, which was not captured by the measured environmental variables.

**Figure 9.**
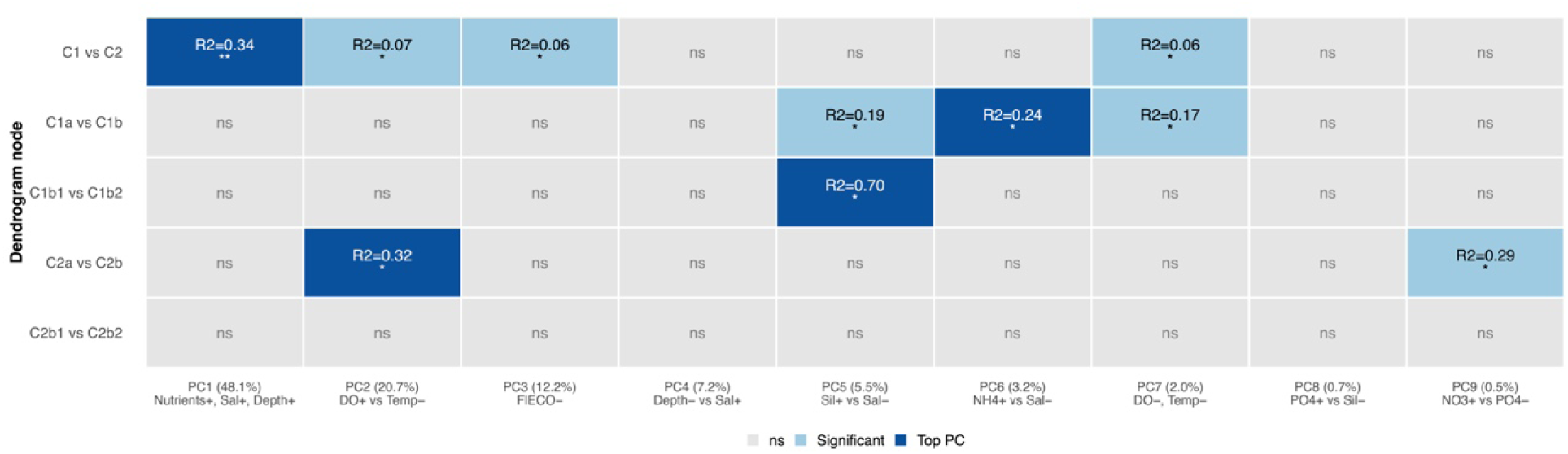
Results of pairwise PERMANOVA on Individual Principal Component (PC) Axes for Each Hierarchical Split Within Division. **C.** Pairwise PERMANOVA was conducted with 999 permutations. P-values were adjusted within each node using the Benjamini-Hochberg method. Each cell represents the PERMANOVA result for a given dendrogram node (rows) and PC axis (columns). The dominant variable loadings for each PC axis are shown alongside, with (+) and (–) indicating the direction of each variable’s contribution. Dark blue cells indicate the top PC axis for each node, defined as the axis with the highest R² among those with significant PERMANOVA (BH-adjusted p < 0.05) and non-significant PERMDISP (p > 0.05). Light blue cells indicate other significant PC axes. Gray cells indicate non-significant results (ns). R² values and significance levels are shown within each significant cell (Significance levels: * p < 0.05, ** p < 0.01, *** p < 0.001).

A dbRDA was conducted to examine the correlations between environmental variables and microbial community composition across the 11 sample clusters (Figure 10). Variance inflation factors (VIF) revealed strong multicollinearity among nutrients (NO_3_, VIF = 31.3; PO_4_, VIF = 22.5; Sil, VIF = 15.7). Variables PO_4_, Sil, NH_4_, and sampling depth were sequentially removed to reduce multicollinearity, resulting in a final set of five variables (temperature, salinity, DO, NO_3_, and FlECO-AFL; all VIF < 3). These variables were scaled to zero mean and unit variance prior to analysis. Seven samples with missing environmental data were excluded, resulting in 174 samples used in this model.

**Figure 10.**
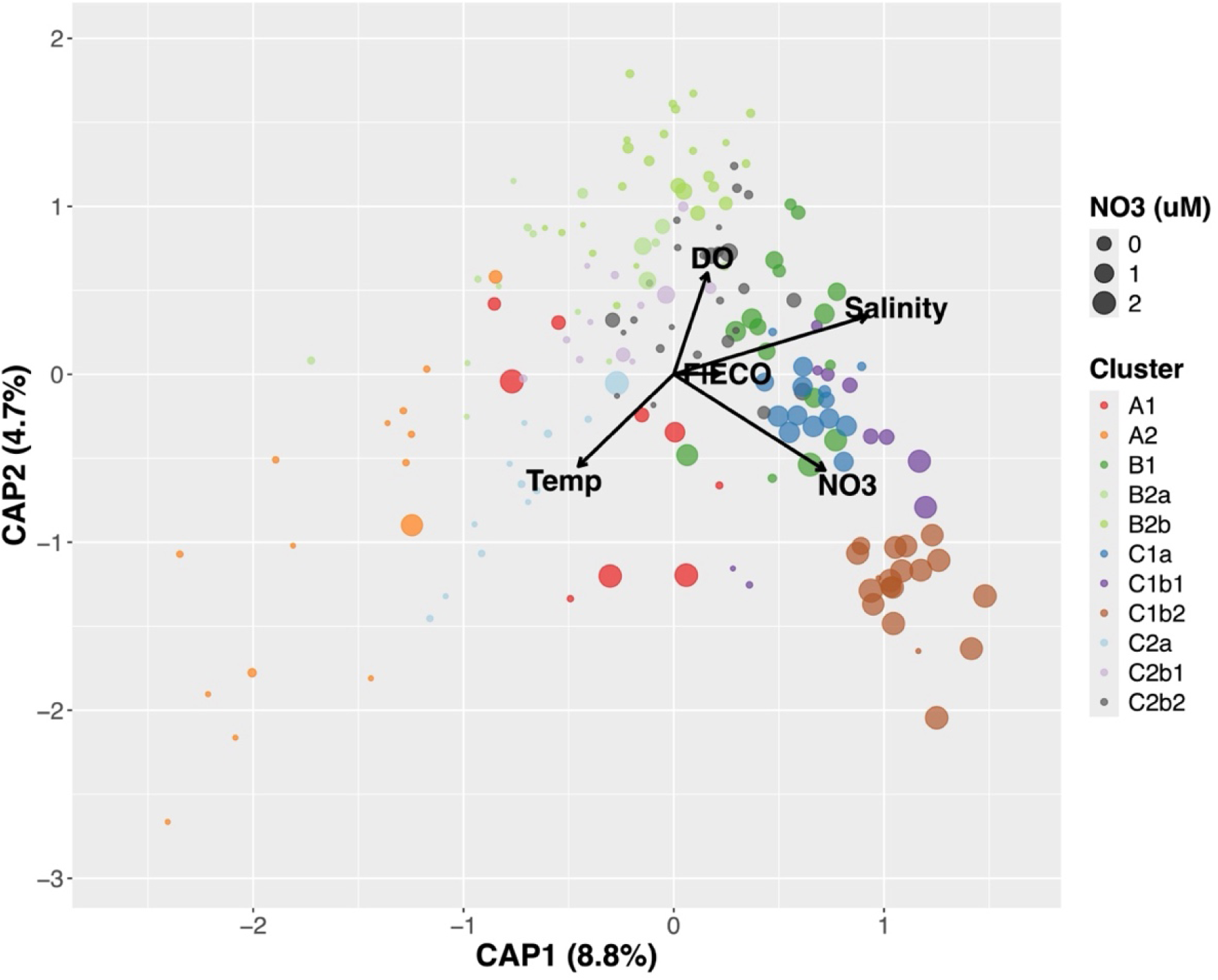
Distance-Based Redundancy Analysis (dbRDA) of the 11 Clusters Constrained by Five Environmental Variables. Temperature, salinity, DO, NO_3_, and FlECO-AFL were used in the dbRDA. Samples are plotted on the first two canonical axes (CAP1, 8.8 %; CAP2, 4.7 %), colored by the 11 sample clusters, with dot size scaled to NO_3_ concentration (µM). Arrows indicate the direction and relative strength of each environmental variable’s contribution to the constrained ordination.

The dbRDA model was statistically significant (F = 7.00, p = 0.001, permutations = 999), with the five environmental variables explaining 14.8 % of the total variation in community composition (adjusted R² = 0.148). Salinity was the strongest contributor to the model (F = 8.44, p = 0.001).

On the CAP ordination, pre-bloom clusters (C1b2, C1a, and B1) were positioned toward high CAP1 values, driven by high salinity and NO_3_, while wCW-influenced post-bloom clusters (A2 and C2a) were positioned at low CAP1 and low CAP2, driven by high temperature and low salinity. Post-bloom clusters in cSW (B2b, C2b2, B2a, and C2b1) occupied high CAP2 positions, associated with high DO. Division A clusters were mostly differentiated by temperature and salinity, while Division B clusters were by DO and salinity. Within Division C, C1 clusters were associated with NO_3_ and fluorescence, while C2 clusters were differentiated by DO and temperature.

A principal component analysis of the nine standardized environmental variables (n = 174 samples) resolved a single dominant axis of variation (Figure 11). PC1 explained 48.1 % and PC2 20.7 % of the total variance (68.8 % cumulative). PC1 was defined by a contrast between the inorganic nutrients, salinity and depth on its positive side (PO_4_, loading 0.46; NO_3_, 0.44; NH_4_, 0.40; silicate, 0.37; salinity, 0.33; depth, 0.31) and temperature on its negative side (–0.28), describing a gradient from warm, fresher, nutrient-poor surface/coastal waters to cold, saline, nutrient-rich, deeper waters. PC2 was mainly driven by dissolved oxygen (0.68) opposed to temperature (–0.51). Samples were distributed as a continuous curved shape along PC1 rather than as discrete groups. The samples curve into an arch rather than falling into separate groups. This arch is a well-known effect that appears when the data are shaped by one strong gradient, so PC2 mostly reflects a curve of that same gradient rather than a truly separate axis. The clusters are therefore not sharply separated environmentally but occupy different positions along one continuous environmental gradient.

**Figure 11.**
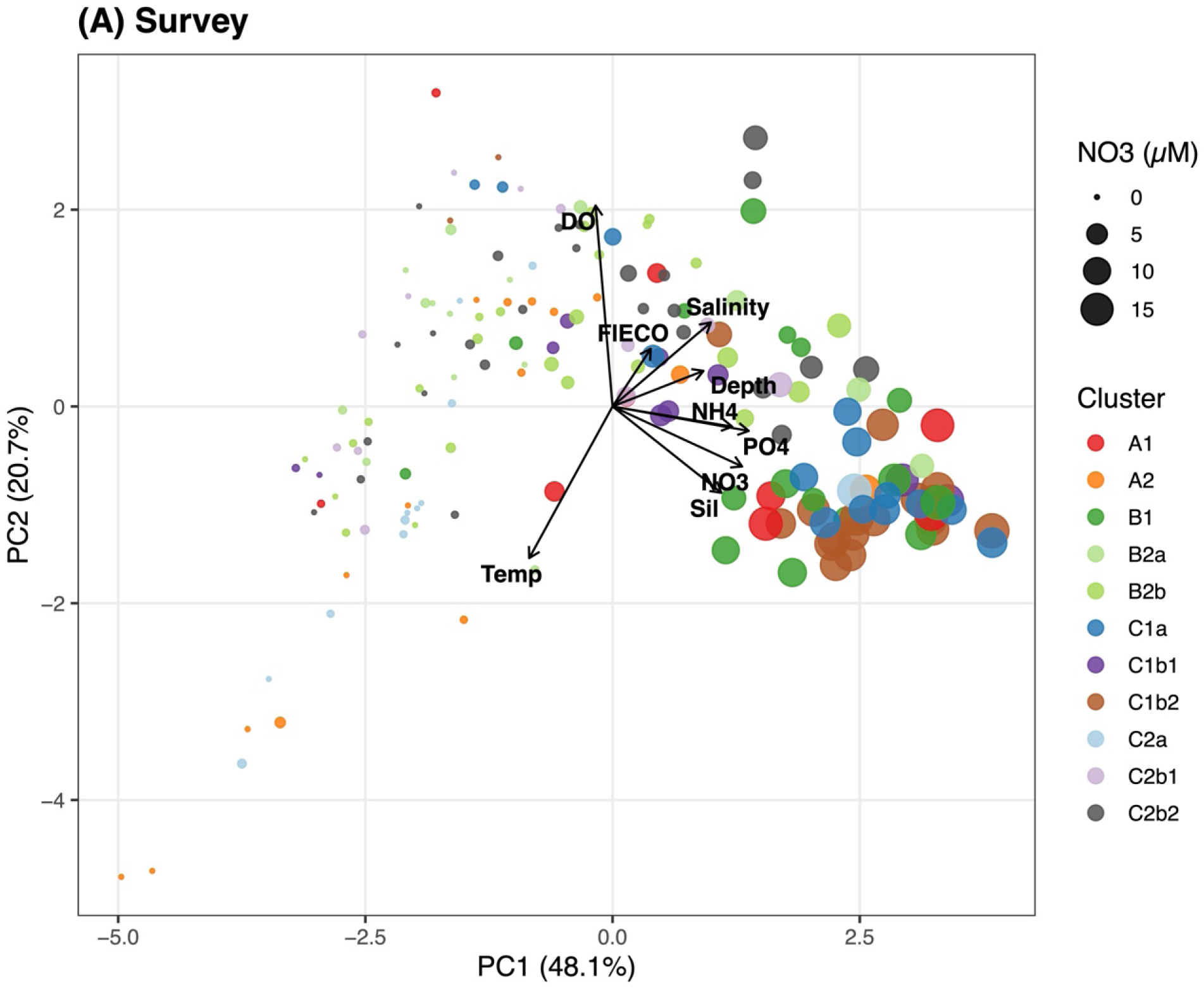
PCA Biplots of the Survey Dataset. Principal component analysis (PCA) of the environmental data of the survey (A) and the process (B) data is shown. Nine variables (temperature, salinity, dissolved oxygen, NO_3_, PO_4_, silicate, NH_4_, fluorescence and depth) were used, with nutrients and fluorescence log-transformed and all variables standardized prior to PCA. Each point is a sample, positioned by its PC1 and PC2, colored by sample clusters, and sized by nitrate concentration (µM). Black arrows show the loadings of the environmental variables. Axes are labelled with the percentage of variance explained (survey: PC1 48.1 %, PC2 20.7 %, n = 174; processing: PC1 48.6 %, PC2 24.8 %, n = 75). PCA was computed separately for each dataset, so scores are not directly comparable between them.

## 4 Discussion

### 4.1 Ecological Characterization of the 11 Clusters and Comparison With Prior Studies

Hierarchical clustering revealed that the surveyed samples harbored distinct microbial communities. Most of the ASVs were classified into three major classes: Bacteroidia, Alphaproteobacteria or Gammaproteobacteria (Figure 12); among each are cosmopolitan free-living taxa and representatives known to respond to algal blooms and contribute to the degradation of algal substrates (Buchan et al., 2014; Bunse & Pinhassi, 2017; Needham & Fuhrman, 2016; Teeling et al., 2012; Teeling et al., 2016).

**Figure 12.**
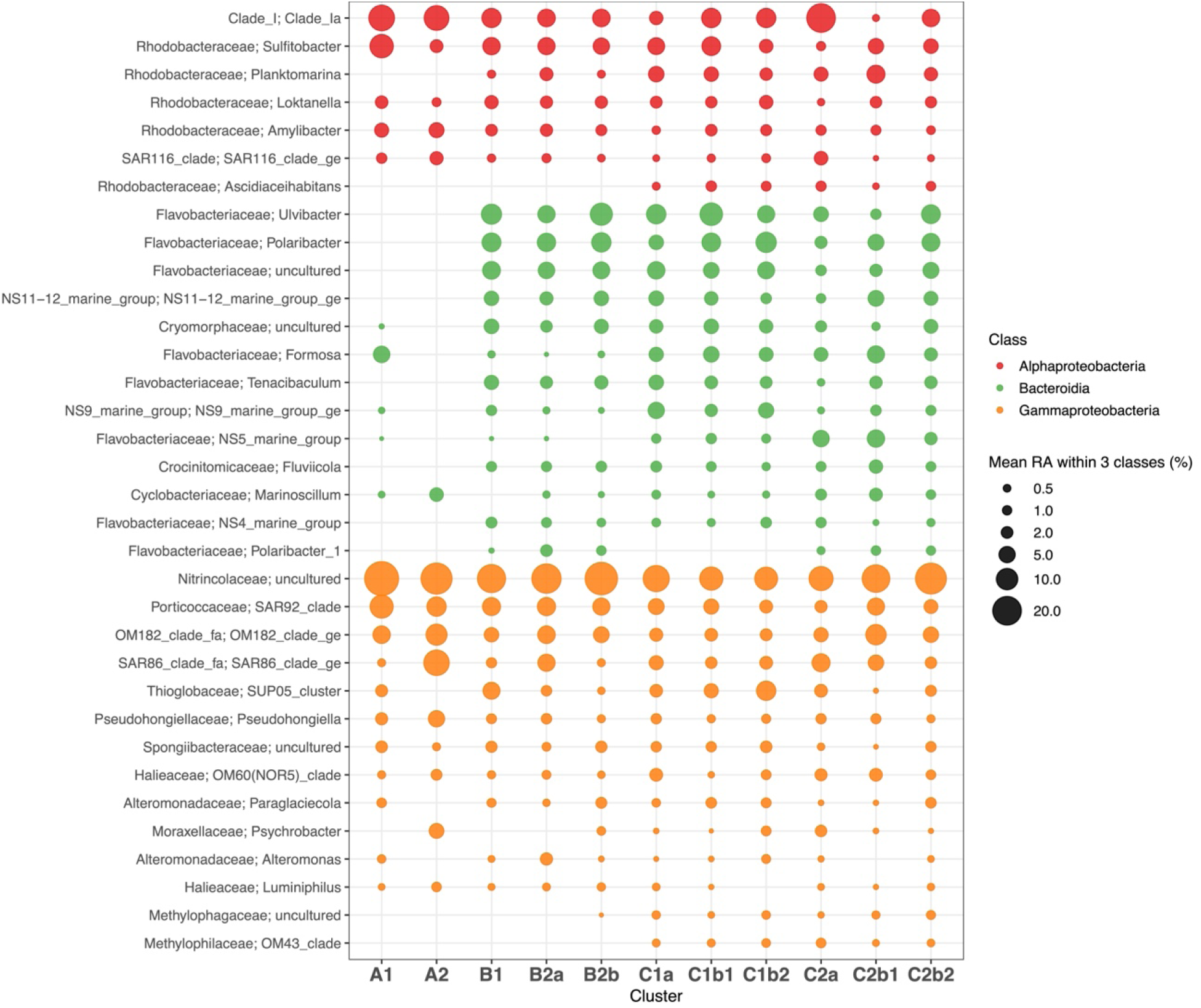
Bubble Plot of Major Genera Within Three Bloom-Responsive Bacterial Classes (Bacteroidia, Gammaproteobacteria, Alphaproteobacteria) Across the 11 Clusters. Rows correspond to genera, columns to sample clusters. Bubble size represents the mean relative abundance (%) of the genus in each cluster among all taxa in the plot, and color represents class. Displayed genera satisfy two objective criteria: occurrence in > 10 % of samples (n = 181) and cumulative within-class relative abundance *≤* 95 % (n = 34 genera: 7 Alphaproteobacteria, 13 Bacteroidia, 14 Gammaproteobacteria).

Within the Alphaproteobacteria, ASVs classified as SAR11 clade Ia and SAR116 clade, and Rhodobacteraceae ASVs such as *Sulfitobacter*, *Amylibacter*, *Planktomarina*, and *Loktanella* were found; these taxa are known to respond to phytoplankton blooms and decomposing organic matter (Bergen et al., 2016; Buchan et al., 2014; Hahnke et al., 2015; Nelson & Carlson, 2012; Sidhu et al., 2023; Takebe et al., 2024; Teeling et al., 2012). Among Gammaproteobacteria, ASVs classified as uncultured Nitrincolaceae, SAR92, OM182, SAR86, SUP05 cluster, and OM60 (NOR5) were abundant, which were also known as degraders of organic matter and association with algal blooms (Cho & Giovannoni, 2004; Dupont et al., 2012; Francis et al., 2021; Morris & Spietz, 2022; Redondo-Río et al., 2024; Ruff et al., 2014; Sakami et al., 2016; Sidhu et al., 2023; Thiele et al., 2022; Wietz et al., 2021; Xue et al., 2021; Yang et al., 2015). For Bacteroidetes, ASVs classified as *Polaribacter*, *Ulvibacter*, uncultured Cryomorphaceae, uncultured Flavobacteriaceae, *Formosa*, *Fluviicola*, *Lutibacter*, *Lutimonas*, and NS marine groups (NS4, 5, 9, 10, and 11-12) were highly abundant; they are representative particle-attached and algae-associated bacteria, contributing to the degradation of algal substrates (Avcı et al., 2020; Bowman, 2020; Choi & Cho, 2006; Ko et al., 2025; Liu et al., 2019; Mann et al., 2013; Meziti et al., 2015; Ngugi & Stingl, 2018; O’Sullivan et al., 2005; Priest et al., 2022; Priest et al., 2023; Thiele et al., 2022; Wietz et al., 2021).

Division A clusters lacked some Bacteroidia ASVs including the eight ASVs missing in Assemblage 1 (Figure 2); two *Ulvibacter* ASVs and one *Polaribacter*, *Fluviicola*, uncultured Cryomorphaceae, *Tenacibaculum*, NS11-12 marine group, and uncultured Flavobacteriaceae. Those ASVs mostly belonged to Bacteroidetes and were highly abundant across samples, and are particularly abundant in sea ice, suggesting recently melted ice may have played a role in structuring communities in the other Divisions. Also, from the boxplots (Figure 7), unique physicochemical characteristics were not found in Division A except for the high variance of temperature. However, this might be the reason for the lack of those ASVs as temperature can be the main factor for bacterial abundance (Korlević et al., 2022), but further investigation is required to answer this question.

In the survey data, total relative abundance of ASVs which belong to Alphaproteobacteria, Gammaproteobacteria, and Bacteroidia accounted for 94.7 %, which is consistent with previous studies conducted in this region (Chen et al., 2008; Lee et al., 2019; Zeng et al., 2012; Zeng et al., 2013a). Remarkably, relative abundance of Gammaproteobacteria ASVs was higher (42.7 %) than Alphaproteobacteria (20.4 %) and Bacteroidia (31.6 %). Bacteroidetes are known for their associations with algal blooms and their contribution to decomposing high molecular weight compounds produced by phytoplankton (Needham & Fuhrman, 2016; Needham et al., 2018). It is also known that the succession of Proteobacteria and Bacteroidetes is often triggered by algal blooms (Bunse & Pinhassi, 2017), and thus, our results were consistent with previous findings, implying the evidence of its association with algal blooms.

Remarkably, Clusters A1 and A2 showed low relative abundance of Bacteroidia (5.5 and 2.9 %, respectively), and this can be ascribed to the factors that caused the eight microorganisms to be missing from these clusters shown in hierarchical analysis (Figure 2). Clusters A1 and C2a showed high abundance of Alphaproteobacteria (34 and 33.1 %, respectively), implying that warm temperature might trigger the abundance of Alphaproteobacteria, as both clusters were largely influenced by warm water masses (wSW and wCW, respectively).

Cluster C2a showed high abundance of SAR11 clade Ia, which is known as one of the most abundant bacteria in oligotrophic marine environment (Giovannoni, 2017), and this can match with the fact that Cluster C2a was the only cluster influenced by nutrient-scarce wCW. Cluster A1 showed high concentration of NO_3_ (Figure 7), so this is not consistent with the possible reason of the abundance of Alphaproteobacteria in Cluster C2a. However, since the mean relative abundance of Bacteroidia ASVs was substantially low in Cluster A1, the proportion of Proteobacteria became extremely high, and thus, we need to interpret this result carefully.

Some pre-bloom clusters such as Clusters C1a and C1b1 showed high abundance of Bacteroidia (40.3 and 40.2 %, respectively), and those clusters were mainly influenced by wSW. However, C1a also showed relatively high abundance of Gammaproteobacteria along with other pre-bloom clusters such as B1 and C1b2. Those three clusters showed balanced proportion of Gammaproteobacteria to Bacteroidia (the difference between the proportions of the two was within 3 %). On the other hand, some post-bloom clusters such as Clusters B2a, B2b, C2b1, and C2b2 showed high abundance of Gammaproteobacteria (46.6, 42.9, 43.3, and 41.7 %, respectively). Clusters B2b and C2b2 were influenced by cSW, while Clusters B2a and C2b1 were in the mixed water mass conditions but were more relatively closely associated with the cSW (Table 2).

Previous studies have surveyed microbial community composition in the Arctic Ocean. For example, a pyrosequencing study conducted in the Chukchi Sea, the Beaufort Sea, and Franklin Bay, reported that Gammaproteobacteria and Alphaproteobacteria shared 22-30 % of all sequences, and the two most abundant taxa (at 97 % similarity) were SAR11 and an uncultured Bacteroidetes (Kirchman et al., 2010). Ghiglione et al. (2012) conducted amplicon sequencing in both Arctic and Antarctic waters, including the Chukchi Sea and found that most taxa in both polar oceans belonged to Gammaproteobacteria, Alphaproteobacteria, and Bacteroidia and several taxa such as SAR11, SAR324, known for their ubiquity and having versatile metabolisms (Malfertheiner et al., 2022). Also, *Polaribacter*, one of the representative bloom-associated bacteria, was an Arctic-specific taxon, underscoring the importance of investigating the link between phytoplankton dynamics and Bacteroidetes succession in the Arctic Ocean. Pedrós-Alió et al. (2015) also reported that microbial communities in temperate regions were dominated by Alphaproteobacteria (approximately 50 %), while microbial communities in the Arctic Ocean contained nearly equal proportions of Alphaproteobacteria and Gammaproteobacteria (approximately 20 % each). Also, the proportion of Bacteroidetes in the Arctic Ocean is higher than the global average (18 % vs. 4 %) (Pedrós-Alió et al., 2015).

Metagenomic analyses also reported the dominance of Alphaproteobacteria, Gammaproteobacteria, and Bacteroidetes in the Arctic Ocean. For example, Cao et al. (2020) analyzed metagenomes from the poles to temperate region and revealed that 90 taxa in phylum level, which was classified from 24,504 taxa, mostly belonged to Alphaproteobacteria, Gammaproteobacteria, and Bacteroidetes showing their high relative abundance in surface and deep-water samples from the Arctic Ocean. Royo-Llonch et al. (2021) analyzed 530 MAGs, which were reconstructed from the Arctic Ocean, and found that more than half of MAGs belonged to Gammaproteobacteria (22.8 %), Bacteroidota (20.9 %), Alphaproteobacteria (11.5 %), and the most abundant 12 MAGs were from unknown species in those three class/phylum and other phyla such as Actinobacteria and SAR324.

The association between bacteria and algal blooms has been studied from temperate regions to the Arctic Ocean. In the North Sea, SAR11 clade taxa were abundant in pre-algal bloom communities, Bacteroidetes such as *Formosa*, *Polaribacter*, and *Ulvibacter* were abundant in during-algal bloom communities, and Gammaproteobacteria such as SAR92 clade dominated post-algal bloom communities (Liu et al., 2013; Sintes et al., 2013; Teeling et al., 2012; Teeling et al., 2016). Those studies were conducted in temperate regions and not in the Arctic Ocean, but our result is consistent with their findings. The study from Franklin Bay found that Bacteroidetes and Alphaproteobacteria taxa such as *Roseobacter* (one of the most abundant marine bacteria and an aerobic anoxygenic photosynthesizer; Brinkhoff et al., 2008), showed the highest abundance in the chlorophyll-a maximum, while Gammaproteobacteria became abundant below the chlorophyll-a maximum (Alonso-Sáez et al., 2008). Microbial community analysis through the northern Barents Sea into the east of Svalbard found that during spring and summer, when sea ice algal bloom occurs, Gammaproteobacteria ASVs such as SAR92, OM60, and Nitrincolaceae and Bacteroidetes ASVs such as *Formosa*, *Polaribacter*, and NS9 marine group became abundant, while SAR11 and some nitrifiers dominated winter samples (Thiele et al., 2023). The metagenomic time-series study, which was conducted in Dease Strait, revealed that 80 % of reads were classified into Alphaproteobacteria (33.2 %), Gammaproteobacteria (28.7 %), and Bacteroidia (21.6 %) (Redondo-Río et al., 2024). Remarkably, Alphaproteobacteria was dominant in the pre-bloom community, while Bacteroidetes increased their abundance in the during-bloom community. Gammaproteobacteria also increased alongside the bloom, though they started to decrease in relative abundance as Bacteroidetes exponentially increased.

In summary, we found that Bacteroidia was relatively more abundant in wSW-influenced pre-bloom clusters, while Gammaproteobacteria was more dominant in cSW-influenced post-bloom clusters. For Alphaproteobacteria, we can assume that they were more abundant in oligotrophic environment, excluding the atypical Cluster A2. Previous studies also illustrated the dynamics of Alphaproteobacteria, Gammaproteobacteria, and Bacteroidetes in the Arctic marine bacterial community responding to phytoplankton blooms and are consistent with the results of our study, suggesting that blooms can be a key driver shaping the seasonal succession of specific bacterial taxa in the Arctic marine ecosystem.

### 4.2 Microbial Community Difference in Depth

In this study, 181 samples were collected from the Bering Sea (n = 72) and the Chukchi Sea (n = 109) from different depths: surface (n = 24), middle (n = 22), and bottom (n = 26) in the Bering Sea; surface (n = 36), middle (n = 42), and bottom (n = 31) in the Chukchi Sea. This differentiated our study from previous studies which collected fewer samples from the same depth in this region (Han et al., 2014; Kirchman et al., 2010; Lee et al., 2019; Zeng et al., 2013a; Zeng et al., 2013b). However, the observed ASV richness within each division was largely independent of sampling depth (Figure 13). In Division A, the median richness was 19.0, 25.0, and 18.5 observed ASVs (bottom, middle, surface); in Division B, 47.5, 50.0, and 44.0; and in Division C, 56.0, 59.5, and 57.0. Kruskal–Wallis tests revealed no significant depth effect for any division after Bonferroni correction (*α* = 0.017): Division A (*χ*² = 3.00, df = 2, p = 0.223), Division B (*χ*² = 2.26, df = 2, p = 0.324), and Division C (*χ*² = 1.19, df = 2, p = 0.551). Consistently, richness did not differ significantly across depths when all samples were pooled (*χ*² = 4.43, df = 2, p = 0.109). This result indicates that the depth at which a sample was collected did not, by itself, predict ASV richness in this region, and that richness was similar across the three depths during the sampling period. This does not mean that vertical gradients were not important. Salinity and nutrient concentrations still varied with depth, but the depth label added no information beyond those variables.

**Figure 13.**
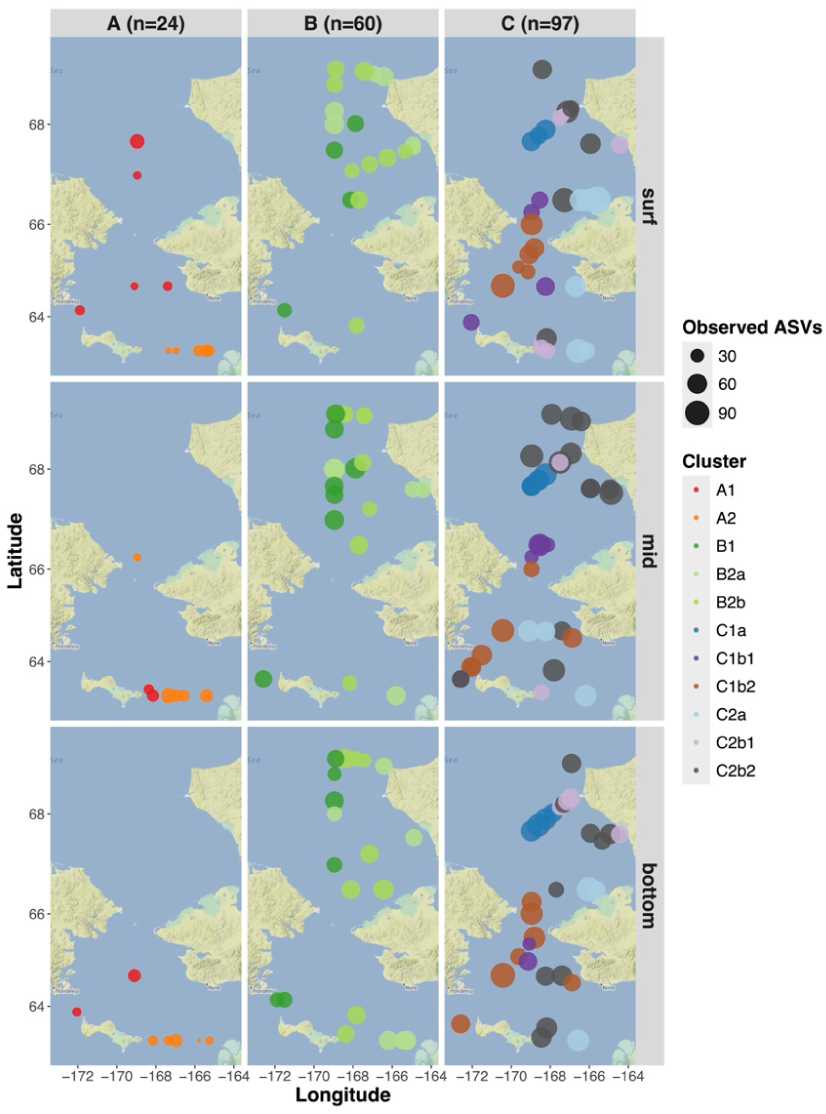
ASV Richness Across Survey Stations, Shown by Depth (Surface, Middle, and Bottom) and Division (A, B, and C). Dot size indicates observed ASV richness (ASVs present within the filtered 258 ASVs). Each dot corresponds to 11 sample clusters.

In both the Bering and Chukchi Seas, very weak thermocline and halocline were observed, whereas no clear clines were evident for the other measured parameters (DO, NO_3_, and chlorophyll-a) (Figure 14). Density anomaly (Δ*σ*_t_) was also calculated to see the strength of stratification in the Bering and Chukchi Seas during June 2017. The Bering Sea exhibited relatively greater vertical stratification than the Chukchi Sea (mean Δ*σ*_t_ = 1.26 vs. 0.43 kg m^−^³, respectively). However, stratification within the Bering Sea was spatially heterogeneous, with 40 % of stations classified as stratified (Δ*σ*_t_ > 1) and another 40 % as well-mixed (Δ*σ*_t_ *≤* 0.5). In contrast, the Chukchi Sea was predominantly well-mixed, with 64 % of stations classified as well-mixed and only 18 % as strongly stratified (mean Δ*σ*_t_ = 0.43 kg m^−^³). The water column in the Bering Sea is stratified during spring when wind weakens and insolation increases (Hunt et al., 2002). However, it is also known that occasional summer mixing happens in this sea (Sambrotto et al., 1986; Whitledge et al., 1986). The Chukchi Sea was single-layered in this study.

**Figure 14.**
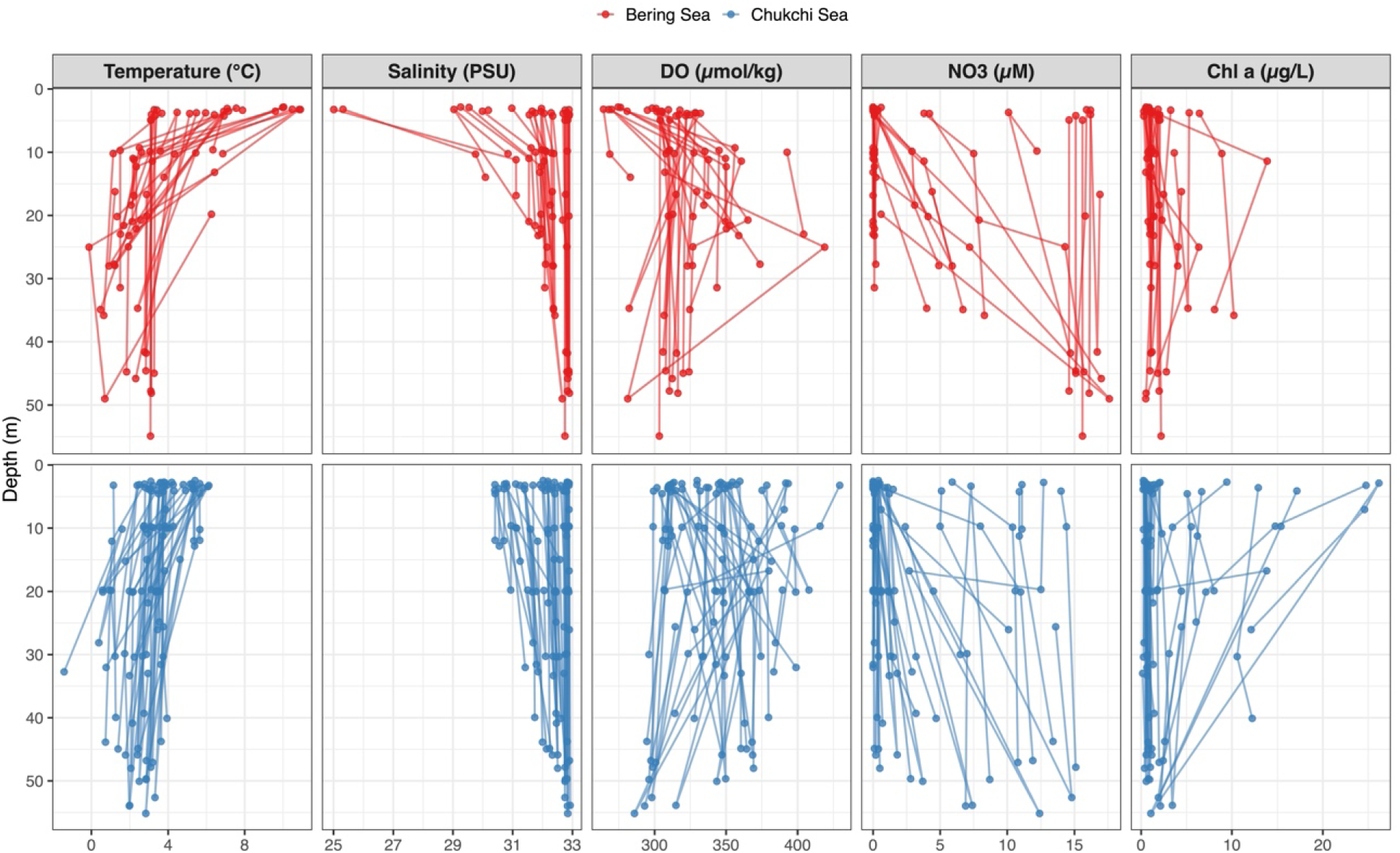
Vertical Profiles of Five Environmental Variables in the Bering and Chukchi Seas. It includes all samples for which the respective variable was measured. Sample sizes differed among variables and between seas. For temperature, salinity, and DO, n = 72 (Bering) and 109 (Chukchi); for NO_3_, n = 71 (Bering) and 103 (Chukchi); and for chlorophyll a, n = 66 (Bering) and 96 (Chukchi). Samples were assigned to the Bering Sea (lat < 65.7°N) or the Chukchi Sea (lat *≥* 65.7°N). Each point represents a single sample plotted at its sampling depth (Y-axis, reversed).

Another possible factor explaining the lack of differences in microbial communities between the surface and deep layers is the transport of microbes to deeper waters with sinking particles. Mestre et al. (2018) examined seawater samples and reported that the dominant prokaryotes on sinking particles in the deep ocean overlapped with those living in surface waters. Also, Boeuf et al. (2019) found that Alphaproteobacteria and Flavobacteria MAGs, which were most prevalent in surface waters, were scarce in deep-sea POM DNA but were relatively abundant in the RNA pool especially in summer, suggesting that these surface-derived bacteria can be exported from the surface to the deep ocean in active-state by attaching to sinking particles, although the authors concluded that sediment trap samples showed distinct surface and deep prokaryotic communities. Those studies compared samples from surface to 4,000 m deep, but given that the Bering and Chukchi Seas are shallow (*≈* 50 m), particle-associated bacteria might be transported quickly from surface waters to bottom waters.

### 4.3 Association of Dominant Bacterial Class with Diatom Bloom

Clustering of the survey samples may have been influenced by biological factors as well as environmental conditions. During the ASGARD 2017 cruise, the high abundance of some diatoms and zooplankton species or groups was reported.

For example, Krause et al. (2021) examined diatom growth and productivity in the northern Bering and Chukchi Seas in June 2017 and reported that diatom biomass (bSiO_2_, µmol Si l^−^¹) was among the highest at station DBO3.8. Consistent with this, Nielsen et al. (2023) found that diatoms dominated the phytoplankton community in the same region in June 2017, as indicated by elevated diatom-associated fatty acid biomarkers (16:1n-7, 16:4n-1, and EPA) and their close association with the June 2017 samples in the Non-Metric Multidimensional Scaling (NMDS) ordination. Lalande et al. (2021) investigated spatial and temporal variation in zooplankton and phytoplankton composition. At DBO3 station in the southern Chukchi Sea, they observed elevated chlorophyll-a fluxes (*≈* 5 mg m^−^² d^−^¹) together with a high abundance of the centric diatoms *Chaetoceros* and *Thalassiosira*, both of which are widespread bloom-forming taxa (De Luca et al., 2019; Park et al., 2016). Because these observations spanned late June to July and coincided with the ASGARD 2017 cruise, they suggest that a pelagic bloom developed around DBO3 during this period. Lekanoff (2020) also conducted clustering analysis of diatom communities in the Bering and Chukchi region. She reported that samples from the ASGARD 2017 cruise were dominated by *Chaetoceros* and *Thalassiosira*, while samples around DBO3 lines were mostly dominated by *Thalassiosira*-rich cluster, which is consistent with the result from Lalande et al. (2021). The association between *Polaribacter* and *Chaetoceros* was reported by microcosm experiments (Takebe et al., 2024), so the results from those studies indicate an association between Bacteroidia and some specific diatom taxa.

Along with diatoms, high abundance of some zooplankton was also reported around DBO stations during the ASGARD 2017 cruise. For instance, *Neocalanus*, a dominant copepod in the northern Pacific Ocean (Conover, 1988; Gonzalez et al., 2025), and polychaete larvae, a globally abundant meroplankton stage (Pamungkas et al., 2019), became highly abundant at DBO2 (the northern Bering Sea) and DBO3 stations during ASGARD 2017 due to the increased inflow of Pacific waters (Lalande et al., 2021).

Based on our results, six clusters contained samples from DBO3 stations: 1 station in Clusters A1 and B2b, 2 stations in Cluster B1, 4 stations in Clusters C1a, C2b1, and C2b2. Especially, all samples in Cluster C1a were collected from DBO3 stations (DBO 3.5–3.8 stations, n = 14). Waffle plot for microbial class composition at eight DBO3 stations (Figure 15A) showed that Gammaproteobacteria, Bacteroidia, and Alphaproteobacteria were highly abundant in each DBO3 station (33-47 % for Gammaproteobacteria, 29-45 % for Bacteroidia, and 17-21 % for Alphaproteobacteria). In addition, chlorophyll-a, FlECO-AFL, and NO_3_ concentration were relatively high around DBO3 stations, especially offshore stations (DBO3.6, 3.7, and 3.8) (Figure 15B), which was consistent with the results from the previous study (Gonzalez et al., 2025) and suggesting that this can be attributed to an active diatom bloom, observed during June 2017 (Krause et al., 2021; Lalande et al., 2021; Lekanoff, 2020; Nielsen et al., 2023).

**Figure 15.**
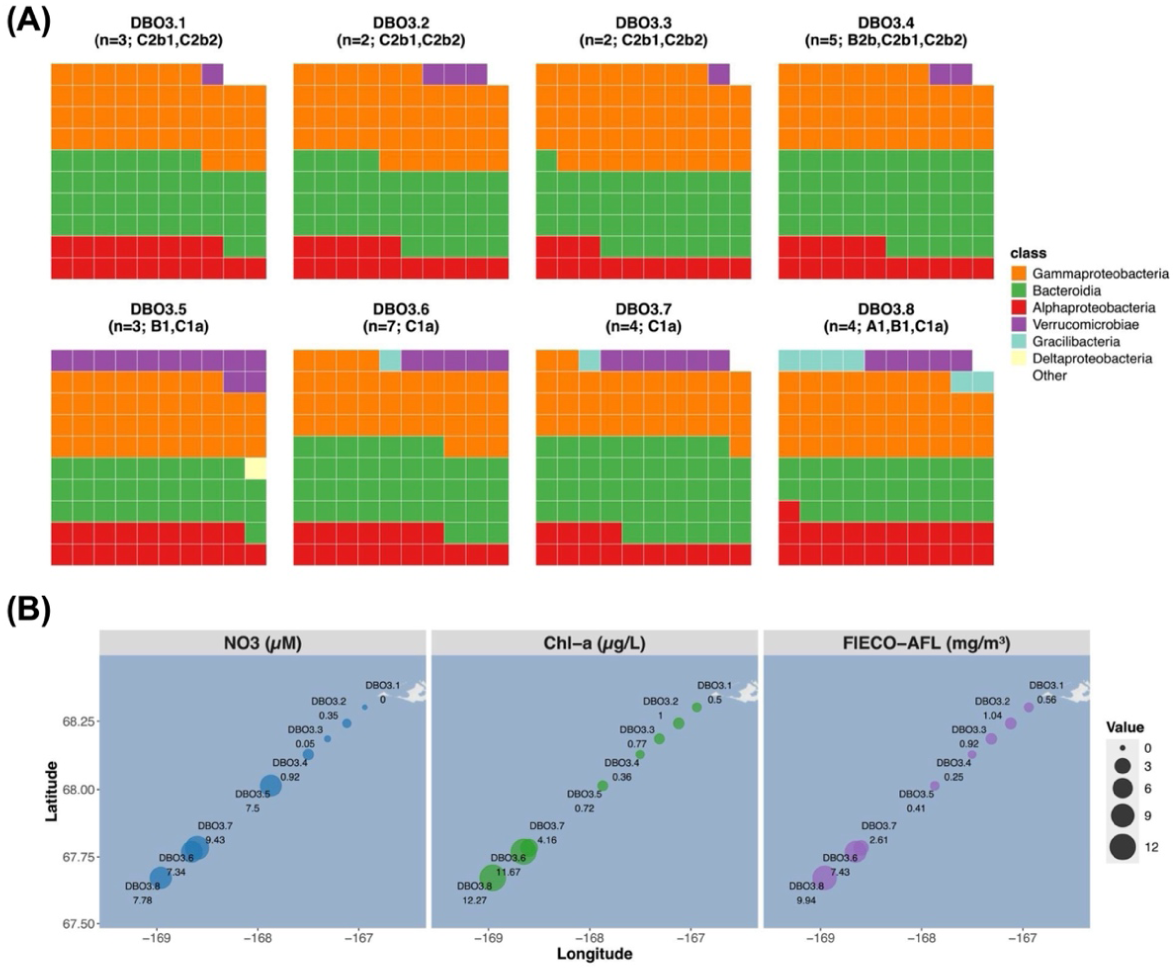
Waffle Plots Showing Class-Level Taxonomic Composition and Map of Nitrate, Chlorophyll-a, and FlECO-AFL Concentrations at DBO3 Stations. *Note*. Samples in six clusters (A1, B1, B2b, C1a, C2b1, and C2b2) were collected in eight stations (DBO3.1-3.8). (A) n in each panel label indicates the number of samples collected at each DBO3 station. Each waffle panel comprises 100 grid cells (10 × 10), with one cell representing 1 % of the within-assemblage relative abundance. Colors indicate taxonomic class; the top 5 classes are shown individually, and the rest are grouped as “Other”; (B) Each point represents each DBO3 station. Point size is proportional to the concentration of each parameter. This map shows the depth-averaged concentration of each parameter.

Compared to other transects, DBO3 showed relatively high abundance of Bacteroidia (Table 3). DBO3 and four other transects (KL, CL, and IL in the Chukchi Sea and CBW in the Bering Sea) showed higher abundance of Gammaproteobacteria and Bacteroidia. Also, the ratio of Alphaproteobacteria: Gammaproteobacteria: Bacteroidia at five transects was approximately 2: 4: 4, suggesting that microbial communities were mainly led by Gammaproteobacteria or Bacteroidia. This pattern further implies that diatom blooms occurred not only along DBO3 but also at the other stations during the ASGARD 2017. In this paper, I focused on DBO3 stations because the pelagic bloom of *Chaetoceros* and *Thalassiosira* was observed around this transect during the ASGARD 2017 cruise (Lalande et al., 2021; Lekanoff, 2020). The bacterial phylum, Bacteroidetes, is known as the association of bloom and the contribution of decomposing complex organic matter such as polysaccharide (Fernández-Gómez et al., 2013; Krüger et al., 2019). *Thalassiosira* is known for its association with Alphaproteobacteria (Mönnich et al., 2020; Pavlovska et al., 2026; Pontiller et al., 2022; Tisserand et al., 2020), Bacteroidetes (Gärdes et al., 2011; Kamalanathan et al., 2019; Mönnich et al., 2020) and Gammaproteobacteria (Pavlovska et al., 2026; Tran et al., 2023). Also, zooplankton such as *Neocalanus* and polychaetes are known to have associations with marine bacteria (De Corte et al., 2014; Kunihiro et al., 2011; Shoemaker & Moisander, 2015; Tan et al., 2015). Taken together, the high abundance of Bacteroidia and Gammaproteobacteria at DBO3 stations and the transects nearby (KL, IL, and CNL) is possibly attributed to the diatom bloom observed in June 2017 and bloom-associated microbial activities.

**Table 3.** Composition of Three Bacterial Classes (Alphaproteobacteria, Gammaproteobacteria, and Bacteroidia) at the 12 Transects in the Survey Data.

| Transect | *n* | Alphaproteobacteria (%) | Gammaproteobacte-ria (%) | Bacteroidia (%) |
|---|---|---|---|---|
| KL | 8 | 12.3 | 43.5 | 40.9 |
| DBO3 | 30 | 16.9 | 38.0 | 40.2 |
| CBW | 16 | 21.6 | 33.7 | 38.3 |
| CL | 20 | 14.0 | 49.2 | 37.6 |
| IL | 13 | 14.9 | 39.6 | 36.6 |
| DBO1 | 4 | 21.2 | 37.2 | 34.3 |
| CPL | 16 | 22.4 | 35.6 | 33.6 |
| DBO2 | 11 | 21.5 | 44.6 | 31.7 |
| CBE | 15 | 20.5 | 42.9 | 31.3 |
| CNL | 22 | 23.2 | 38.6 | 30.4 |
| CB | 2 | 14.7 | 43.5 | 27.1 |
| SLY | 24 | 26.4 | 58.4 | 5.3 |

*Note.* The number of samples is shown in the *n* column. Each proportion is the median relative abundance of the class.

### 4.4 Influence of Water Mass on Microbial Communities

The objective of this study is to investigate the influence of water mass on microbial communities in the Bering and Chukchi Seas. As Table 2 showed, the 11 sample clusters were influenced by different water masses. Clusters B1, C1a, and C1b2, which were wSW-influenced and pre-bloom clusters, showed correlations with NO_3_ and salinity on dbRDA (Figure 10). Also, those three clusters showed higher NO_3_ concentration than other clusters (median: 15.1, 10.5, and 7.7; mean ± SD: 13.17 ± 5.05, 8.62 ± 4.39, and 7.97 ± 5 in Clusters C1b2, C1a, and B1, respectively, Figure 7) and contained balanced proportions of relative abundance of major bacterial classes (approximately 2: 4: 4, Alphaproteobacteria: Gammaproteobacteria: Bacteroidia). However, samples in those clusters were collected from different geographical locations (B1 in the Chukchi Sea including DBO3 stations, C1a in DBO3 stations, and C1b2 in the Bering Sea), across a latitudinal range spanning more than 300 km. This result suggests that these communities were transported with Pacific water flowing northward through the Bering Strait or transported by water that formed preferable environmental conditions for those clusters.

In addition, the six assemblages differed in where they were most abundant, each reaching its highest relative abundance in a different part of the study area (Figure 3). This shows that the communities are spatially structured. However, location and water mass are not independent here. Water flows northward across the shelf, so where a sample was collected also reflects how long its water has been modified by biological activity. Therefore, the spatial pattern reflects both water-mass origin and bloom progression, and the two cannot be separated with these data.

Post-bloom clusters showed the influence of water mass. For example, A2 and C2a, both wCW-influenced clusters (Table 2), showed low loadings on CAP1 and CAP2 but showed correlations with temperature on dbRDA (Figure 10). C2a had the highest temperature (median: 5.4; mean ± SD: 5.68 ± 2.69, Figure 7), while A2 showed the third highest median and the second highest mean ± SD in temperature (median: 3.75; mean ± SD: 5.2 ± 3.68, Figure 7). In addition, compared to other clusters, the relative abundance of Alphaproteobacteria was higher in these clusters (33.1 % in C2a and 25.3 % in A2) though most clusters showed less than 20 % of their abundance. Geographically, sampling locations of Clusters A2 and C2a are different as samples in A2 and C2a were mostly collected from SLY Transect and Alaskan coasts respectively. wCW is nutrient-scarce water, and both A2 and C2a showed the lowest median (0.1 µM; mean ± SD 1.19 ± 3.52 and 1.48 ± 4.86 µM, respectively) of NO_3_ concentration. Also, the most dominant Alphaproteobacteria ASV in Cluster A2 and C2a was SAR11 clade Ia, the representative marine bacteria in oligotrophic environment (Giovannoni, 2017), suggesting that oligotrophic condition in wCW favored the microbial communities in those post-bloom clusters, which were led by Alphaproteobacteria.

The other four post-bloom clusters also represented distinct characteristics. For instance, Clusters B2a and C2b1, which were influenced by both cSW and wSW (Table 2), showed high DO concentration (median: 349.58 and 347.39; mean ± SD: 349.14 ± 32.5 and 350.32 ± 34.51, respectively, Figure 7) and had correlations with DO concentration (Figure 10). Clusters B2b and C2b2, which were cSW-influenced clusters (Table 2), also showed a similar pattern with DO (Figure 7 and Figure 10) and lower nutrient concentrations (median: < 0.5 µM in those four clusters, Figure 7). Also, microbial communities in these four clusters were led by Gammaproteobacteria (46.6 % in B2a, 43.3 % in C2b1, 42.9 % in B2b, and 41.7 % in C2b2). Mostly, Gammaproteobacteria is known as increasing their abundance in post-bloom conditions (Liu et al., 2013; Sintes et al., 2013; Teeling et al., 2012; Teeling et al., 2016), and dominant ASVs in these four clusters (Figure 12) were uncultured Nitrincolaceae, SAR92, OM182, and SAR86, which were known as bloom-responding carbon degraders (Sidhu et al., 2023; Thiele et al., 2022; Wietz et al., 2021; Yang et al., 2015). Sampling locations of Clusters B2b, C2b1, and C2b2 were mainly from the Chukchi Sea (B2b from CL and IL, C2b1 from DBO3, and C2b2 from KL, IL, and DBO3). On the other hand, most samples in Cluster B2a were collected from both the northern Chukchi Sea (CNL, 29.4 %) and the southern Bering Sea (SLY, 23.5 %). Although samples in Cluster C2b1 were largely collected from DBO3 stations (50 %), it also contained samples from the southern Bering Sea (SLY, 8.3 %; CBE, 25.0 %). This can be the reason why Clusters B2a and C2b1 showed similarities in water mass conditions and microbial community composition.

Those results support our interpretation that microbial community composition across the Bering and Chukchi Seas is structured by water mass origin and bloom-stage progression along with geographic distance. However, we must remember the fact that sea water temperature in this region anomalously increased and experienced early-ice retreat due to the heat wave during June 2017 (Baker et al., 2020; Danielson et al., 2020; Walsh et al., 2018). A mooring study also reported that heat transport from the Pacific Ocean to the Canada Basin increased 1.5 times over 20 years (Itoh et al., 2026). Danielson et al. (2020) analyzed water mass distribution in the Bering and Chukchi Seas and showed that before 2014, cSW was prevalent in the upper water column through this region due to seasonal warming and sea ice melt. In this paper, we investigated the proportion of influential water mass on each sample cluster (Table 2) and found that primary water masses of most sample clusters were either wSW or cSW. Given the impact of the heat wave, sample clusters classified as having wSW as the primary water mass might have been cSW-influenced clusters in fact.

The Bering and Chukchi regions are known for their high productivity, but it is also known that they are vulnerable to sea ice loss. Ballinger & Overland (2022) reported that dramatic loss of sea ice in the Bering and Chukchi regions and shifted atmospheric patterns led the Alaskan Arctic to enter a new climate regime in 2017-2021, and it may cause the unprecedented extreme events such as delaying spring blooms and disrupting marine food webs. Furthermore, microbial communities in the Arctic Ocean are influenced by sea ice loss. For example, Comeau et al. (2011) analyzed post-bloom communities in the subsurface chlorophyll maximum layer in the Beaufort Sea before and after the loss of MYI in 2007 and found that the abundance of Bacteroidetes dramatically decreased after 2007, and more than half of this reduction was attributed to *Polaribacter* and unidentified Flavobacteriaceae. On the other hand, the relative abundance of Alphaproteobacteria was consistently high, and Gammaproteobacteria did not change their abundance much. Kraemer et al. (2024) also conducted time-series analyses of microbial community composition in the Canada Basin and reported that the abundance and diversity of Bacteroidetes dramatically decreased after sea ice loss, and the majority of decreasing ASVs were members of the Bacteroidetes and Proteobacteria. These studies collectively demonstrated the significant impact of sea ice dynamics on the composition and succession of Bacteroidetes in Arctic marine microbial communities. In this paper, we found that wSW-influenced clusters and cSW-influenced clusters had different microbial community compositions (Figure 2; Table 2; Figure 12), implying that future environmental shift probably alters the microbial compositional change in this region, and thus, we need to monitor microbial communities as a sentinel of environmental change.

## Data availability

Raw 16S rRNA gene amplicon reads from the ASGARD 2017 samples are available in the NCBI Sequence Read Archive under BioProject PRJNA421293 (SRA study SRP126234; https://www.ncbi.nlm.nih.gov/bioproject/PRJNA421293). All R scripts and outputs are publicly available at https://github.com/ReiSHIBUE/ASGARD2017_analysis.

## Author contributions

R.S. analyzed the data and wrote the manuscript. R.E.C. supervised the study.

## Acknowledgements

We thank Rachel M. Lekanoff, who participated in the ASGARD 2017 cruise, collected the DNA and environmental samples, processed them at the UAF Water and Environmental Research Center and the UAF DNA Core Lab, and provided the microbiological, hydrographic and environmental dataset.

